# Glycosylated diterpenes associate with early containment of *Fusarium culmorum* infection across wheat (*Triticum aestivum* L.) genotypes under field conditions

**DOI:** 10.64898/2025.12.02.691979

**Authors:** Stefan A. Pieczonka, Fabian Dick, Melissa Bentele, Ludwig Ramgraber, Lukas Prey, Erwin Kupczyk, Johannes Seidl-Schulz, Anja Hanemann, Patrick O. Noack, Stefan Asam, Philippe Schmitt-Kopplin, Michael Rychlik

## Abstract

Wheat (*Triticum aestivum* L.) is severely affected by head blight, a destructive disease caused primarily by *Fusarium* species, which reduces yield and contaminates grains with trichothecene mycotoxins that compromise food and feed safety worldwide. Mechanistic insights into host–pathogen interactions have largely emerged from controlled experimental settings, providing conceptual foundations under conditions of constrained genetic and environmental complexity. Resolving consistent, generalizable molecular patterns across diverse wheat genotypes within a large-scale, heterogeneous field study remains technically and methodologically challenging. Addressing this gap, we integrated quantitative mycotoxin profiling with untargeted metabolomics in a field experiment comprising 105 wheat genotypes deliberately inoculated with *F. culmorum*, with the goal of identifying molecular signatures of infection and containment under realistic agronomic conditions. Quantified deoxynivalenol (DON) concentrations provided a robust, continuous measure of infection intensity, enabling metabolite profiles to be related directly to pathogen activity beyond the limitations of symptom-based visual scoring. Consistent with established *Fusarium* trichothecene biosynthesis, sesquiterpene-derived metabolites closely tracked toxin accumulation, thereby recapitulating infection-associated metabolic patterns across the large-scale field study. Conversely, glycosylated diterpene conjugates were elevated under low toxin accumulation, linking their abundance to contained *Fusarium* pathogen activity and highlighting a largely underexplored aspect of wheat metabolism in host–pathogen interactions. Elucidating their biosynthetic origin, enzymatic interconversion, and regulatory context will be key to defining potential diterpene-associated defense processes across wheat genotypes.

## Introduction

Wheat (*Triticum aestivum* L.) is one of the world’s most important staple crops and a cornerstone of global food security. However, its productivity is severely threatened by Fusarium head blight (FHB, also known as wheat scab), a destructive disease of small-grain cereals. FHB not only reduces yield and grain quality but also results in the accumulation of toxic secondary metabolites (mycotoxins), which compromise food and feed safety, causing substantial economic losses worldwide (1). The disease is primarily caused by *Fusarium graminearum, F. culmorum*, and *F. avenaceum* species that differ in regional prevalence but share the ability to produce harmful mycotoxins. Due to this dual impact on crop performance and safety, *Fusarium* resistance is, alongside drought tolerance, growth stability, and yield, a key breeding objective (2). Infections typically develop under warm (above 16 °C), humid conditions and are most likely between mid-ear emergence (BBCH 50–58) (3) and the end of flowering (BBCH 60–69) (4). Residues of previous cereal crops serve as major reservoirs of inoculum, while characteristic symptoms include the premature bleaching of spikelets, which can progress to complete colonization of ears within 10–14 days (5). Resistance is commonly assessed by visual phenotyping in early summer; however, this approach is labour-intensive and subject to environmental variability and the subjectivity of expert evaluation.

From a consumer perspective, mycotoxin contamination is of greater concern than visible disease severity. Deoxynivalenol (DON), the principal toxin of *F. culmorum*, exerts acute and chronic toxicity and is subject to strict legal limits in many countries (6). Although symptom severity generally is associated with DON accumulation (7), field scoring provides only an indirect estimate. Modern approaches therefore increasingly employ unmanned aerial vehicles (UAVs), whose high-resolution imaging combined with predictive models enables more efficient and consistent evaluation of FHB severity (8) and mycotoxin load. Although such novel phenotyping tools will improve the efficiency of field-based infestation assessment (9, 10), the direct monitoring of mycotoxin concentrations in grain remains indispensable for food and feed safety. A wide range of analytical methods is available, including ELISA, HPLC‐UV, HPLC‐FLD, GC‐MS, and, most commonly, LC–MS/MS. ELISA serves as a rapid screening approach for single toxin groups (11). HPLC-coupled UV and fluorescence detection has traditionally been used for quantification, still remain valuable for specific analytes, but is increasingly outperformed by more sensitive alternatives. Mass spectrometry-based methods, especially LC–MS/MS, have become standard in modern applications (12–15), reporting substantial year-to-year variation in DON concentrations in wheat field studies, with levels reaching up to 10 mg kg^-1^ in both *Fusarium*-inoculated and naturally infected crops depending on environmental conditions (16–18). Their ability to simultaneously detect structurally diverse toxins and to achieve accurate quantification in complex matrices—particularly when combined with stable isotope dilution analysis—provides superior sensitivity, precision, and robustness (11). For instance, an LC–MS/MS method recently established in our group for the simultaneous quantification of *Alternaria* and *Fusarium* toxins in cereal samples illustrates how isotope-labelled standards, by compensating for matrix effects and work-up losses, enable robust and reliable quantification (19).

In addition to the targeted quantification of mycotoxins, omics-based studies offer insights into the molecular processes underlying fungal infection, plant defense, and resistance mechanisms. The infection biology of *F. graminearum* has been extensively investigated and is known to rely on signaling pathways (MAPK, cAMP–PKA), transcriptional regulators (Tri6, Tri10), and effector proteins such as Osp24 and FgL1, which collectively coordinate DON biosynthesis, suppress host defenses, and promote fungal spread (20). DON itself acts as a central virulence factor by suppressing host defense responses (21), modulating programmed cell death (22), and facilitating fungal expansion within the wheat head (23). On the host side, detoxification mechanisms such as glucoside or glutathione conjugation (24) and vacuolar sequestration have been described. Further defense responses include the restriction of pathogen advancement (25, 26) and reinforcement of structural barriers through antimicrobial and lignin-related hydroxycinnamic acid derivatives (27). Wheat-characteristic phytoanticipins such as benzoxazinoids (28, 29) likewise contribute to preformed chemical defense. Despite these advances, studies on the metabolic interface between host and pathogen—the level most directly linked to the phenotype—often remain restricted to a few cultivars grown under narrowly defined conditions, limiting their general explanatory power (30, 31). Recent findings (32) further highlight the complexity and cultivar-specific nature of FHB responses. However, ongoing developments in metabolomics increasingly enable large-scale, comparative analyses that can detect shared molecular patterns even among unknown compounds.

In this field study across 105 *Triticum aestivum* L. genotypes—comprising 72 breeding lines, 24 of which are commercial varieties—grown under naturally heterogeneous field conditions, we combined accurate quantification of *F. culmorum* toxins with untargeted metabolomics to move beyond the limitations of visual FHB scoring. By correlating toxin load with the metabolome of infected grains across genetic and nutritional backgrounds, we aim to identify common metabolic signatures reflecting infection progression and host response. Integrating objective, quantitative, and continuous toxin measures provides a framework to investigate infestation-induced metabolic changes under realistic field variability. By linking molecular responses to precisely quantified toxin burdens rather than subjective and distinct visual classifications, this approach is proposed to enable the identification of robust plant–pathogen interaction patterns that transcend specific cultivars or constrained laboratory-scale experimental settings.

## Methodology

### Field study

Wheat samples were obtained from a winter wheat field trial conducted in 2022 near Herzogenaurach (southeastern Germany), where *F. culmorum* inoculation, standard agronomic practice, and field-based FHB severity scoring were performed under drought-affected conditions following established breeding evaluation protocols detailed in Rößle *et al*. (10). The field study was part of the AutoDGB project (project number 2818407A18) funded by the German Federal Office for Agriculture and Food (Bundesanstalt für Landwirtschaft und Ernährung) on optical field analysis of wheat, with related publications detailing non-invasive FHB estimation using RGB imaging, multispectral and RGB-based yield phenotyping, UAV-based grain yield prediction, and spectral plot-level aggregation approaches (9, 10, 33–35). Details on soil and weather conditions are provided in Rößle *et al*. (10). The germplasm comprised preselected material from F5 generation onward and double-haploid lines, representing a spectrum of known FHB susceptibilities while excluding genotypes with extreme morphological or phenological traits. No fungicide was applied, and *F. culmorum* spores were inoculated twice (30th May and 1st June 2022), corresponding to early and late milk ripeness stages, at concentrations of 10^5^ spores L^-1^ in 600 L ha^-1^.

FHB severity was rated on a 1–9 scale following the scale of the German Federal Plant Variety Office (36), corresponding to increasing infection from 1 (no symptoms) to 9 (61–100% of ears infected), defined as follows: 1 (no infection), 2 (0–2% infection), 3 (2–5% infection), 4 (5–8% infection), 5 (8–14% infection), 6 (14–22% infection), 7 (22–37% infection), 8 (37–61% infection) and 9 (61–100% infection). Samples were visually evaluated twice in June during milk ripeness, when symptoms are most reliably distinguished in standard field practice, and collected in August at physiological maturity to capture the final toxin burden and metabolomic state of the harvested grain. An overview of genotype variability and observed infection severity is provided in **Supplementary Table S1**.

### Sampling strategy

Sampling aimed to capture a representative range of infestation severity. In total, 145 plots representing 105 genotypes—comprising 72 breeding lines, 24 of which are commercial varieties—were selected to capture the full range of visible FHB symptoms observed in the field. This design balanced statistical representativeness for large-scale data analysis with the precision required for toxin quantification. Ears were manually harvested from each plot, grains (100–150 g per sample) were collected for analysis, stored, and transported at room temperature before being ground within one week of arrival.

### Mycotoxin quantification

Mycotoxin quantification was performed using our previously developed and validated targeted LC–MS/MS method employing a LCMS-8050 triple quadrupole mass spectrometer (Shimadzu, Kyoto, Japan) (19). Details on mass-spectrometric parameters and multiple-reaction-monitoring transitions are provided therein and summarized in **Supplementary Table S2**. The method enables quantification of 23 mycotoxins in cereals. Following qualitative assessment of all detected compounds, our analysis focused on DON, the principal toxin of *F. culmorum*, and its derivatives 3-acetyl-DON (3-AcDON), a metabolic precursor, and DON-3-glucoside (DON-3-G), a plant-derived detoxification conjugate. As no fungicides were applied, natural *Alternaria* infestation was evaluated through quantification of its principal toxin, tenuazonic acid (TeA).

Quantification of the mycotoxins DON, 3-AcDON, and TeA was carried out using stable isotope-labelled standards of the respective compounds. DON-3-G was quantified via a matrix-matched calibration curve, relating its signal to the ratio between DON-3-G and [^13^C_15_]-DON, thereby retaining the advantages of stable-isotope dilution. For DON-3-G, a three-point calibration curve (10, 50, and 100 µg kg^-1^) was established and analyzed in duplicate with each batch of samples. The correlation coefficients exceeded 0.99 in all cases, and linearity was confirmed using Mandel’s fitting test (37). Limits of detection (LODs) and limits of quantification (LOQs) were 0.32 and 0.87 µg kg^-1^ (DON), 1.96 and 7.42 µg kg^-1^ (DON-3-G), 0.28 and 0.81 µg kg^-1^ (3-AcDON), and 0.14 and 0.50 µg kg^-1^ (TeA), respectively, as reported in Dick *et al*. (19).

### Sample extract preparation

A QuEChERS (38) cleanup was employed as a unified sample preparation approach for both targeted mycotoxin quantification (19) and untargeted metabolomics analysis. To quantify toxin content, a representative portion of each sample (80–150 g) was pooled and ground to fine wholegrain flour using a laboratory mill (Retsch, Haan, Germany). One gram of the homogenized material was used for analysis and spiked with internal standards: 50 µL of 1 µg mL^-1^ [^13^C_6_,^15^N]-TeA (39), 50 µL of 1 µg mL^-1^ [^13^C_15_]-DON (Biopure, Tulln, Austria), 20 µL of 0.1 µg mL^-1^ [^13^C_17_]-3-AcDON (Biopure, Tulln, Austria), and 50 µL of 0.01 µg mL^-1^ [^15^N_3_]-ENN A_1_ (40). Extraction followed the protocol optimized and established recently (19). Sample workup was performed in duplicate, resulting in 290 samples of 145 individual plots. When the coefficient of variation for a given toxin exceeded 25%, the analysis was repeated to ensure accuracy. Elevated variation was attributed to potential inconsistencies during extraction or measurement, or to sample inhomogeneity (41).

### Mycotoxin and FHB correlation

Residual diagnostics of linear models indicated pronounced heteroscedasticity of DON concentrations across FHB severity classes and deviations from normality. Therefore, differences in DON concentrations between visual severity classes were assessed using Welch’s ANOVA, which does not assume equal variances. Post-hoc pairwise comparisons were performed using the Games–Howell test. Adjusted p-values were converted into Compact Letter Displays (CLD) to summarize statistically distinct severity groups. All analyses were conducted in R (version 4.5.0) using the rstatix and multcompView packages and are summarized in **Supplementary Table S3**.

### Metabolomics measurement and data processing

The QuEChERS extracts were analyzed on a Sciex X500R QTOF UHPLC–ToF–MS system (Sciex, Framingham, MA, USA) equipped with an ACQUITY UPLC BEH C18 column (Waters, Milford, MA, USA). Data were acquired in both positive and negative electrospray ionization (ESI) modes using SWATH acquisition (42). The UPLC gradient (**Supplementary Table S4**) and instrument parameters (**Supplementary Table S5**) followed previously published methods (43). A pooled QC sample, prepared by combining equal aliquots of all extracts, was injected in duplicate every 15 runs to monitor and normalize signal stability throughout the batch. As negative ionization yielded only limited complementary information, subsequent data processing and interpretation focused on the ESI-positive mode.

The resulting.wiff2 profile data files were imported into the MS-DIAL 4.2 open-source data processing environment (44). Detailed settings for data collection, peak detection, spectrum deconvolution, adduct ion identification, alignment, and data export are provided in **Supplementary Table S6**. In brief, measurement parameters were configured according to the MS acquisition method. An exclusion mass list was applied to remove masses corresponding to the calibrant mix and common solvent contaminants, minimizing noise features. SWATH spectrum deconvolution utilized the software’s intrinsic machine learning-based approach to resolve overlapping fragment ion signals and assign them to precursor ions based on retention time and *m/z* profiles. For peak alignment, QC measurements served as reference files, with retention time correction, MS1 adjustments, peak count filtering, and abundance-based filtering (2 of 2 replicates) within sample groups applied. A blank correction was implemented. Following alignment table generation, batch effects were corrected using LOWESS regression to harmonize feature intensities across QC measurements. The final feature matrix was exported in.mgf format for further analysis using SIRIUS 5.8.2 (45).

### Metabolomics statistics and data interpretation

A principal component analysis (PCA) was performed using the MS-DIAL feature list to evaluate overall metabolite diversity and identify underlying correlations in an unsupervised manner. Prior to statistical modelling, chromatographic feature intensities (peak heights) were zero-filled and subsequently normalized using Z-scores (**Supplementary Table S6**). A supervised orthogonal partial least squares (OPLS) analysis was performed to correlate metabolomics data with toxin levels, using quantified DON concentration as the latent Y-variable. Features with a Variable Importance in Projection (VIP) score ≥2.5 were considered the most significant positive or negative contributors. Model quality was evaluated based on R^2^Y (goodness of fit) and Q^2^ (predictive ability) values (46). The correlation strength of the most significant features was further examined using linear correlation and regression analyses (*α* = 0.05). To characterize the molecular structures of correlated metabolites, the exported.mgf peak list was analyzed in the SIRIUS 5.8.2 software environment (45). Tandem MS data were used to derive molecular formulas via fragmentation trees (47), assign compound classes using CANOPUS (48), and predict candidate structures through CSI:FingerID (49). The databases consulted and detailed parameter settings are provided in **Supplementary Table S7**. Identification confidence levels followed the framework of Schymanski *et al*. (50), integrating database matches, diagnostic fragment ions, and in silico fragmentation predictions.

## Results and Discussion

### Field study and plot rating

FHB severity was visually assessed by whitening symptoms on wheat ears during the milk stage. Each plot was assessed twice as a technical repeat, with evaluations carried out in early June 2022, yielding scores from 1 to 8 (**Figure 1**): 1 (29 plots), 2 (41), 3 (52), 4 (57), 5 (57), 6 (35), 7 (15), 8 (4), and 9 (0). The evaluations confirmed successful inoculation, and the range of visual severities reflected the expected variation in susceptibility among varieties as well as natural variation of individual plots. Differences between the two evaluation dates were minor and primarily reflected scoring uncertainty and differences in developmental stage and field conditions.

**Figure 1.**
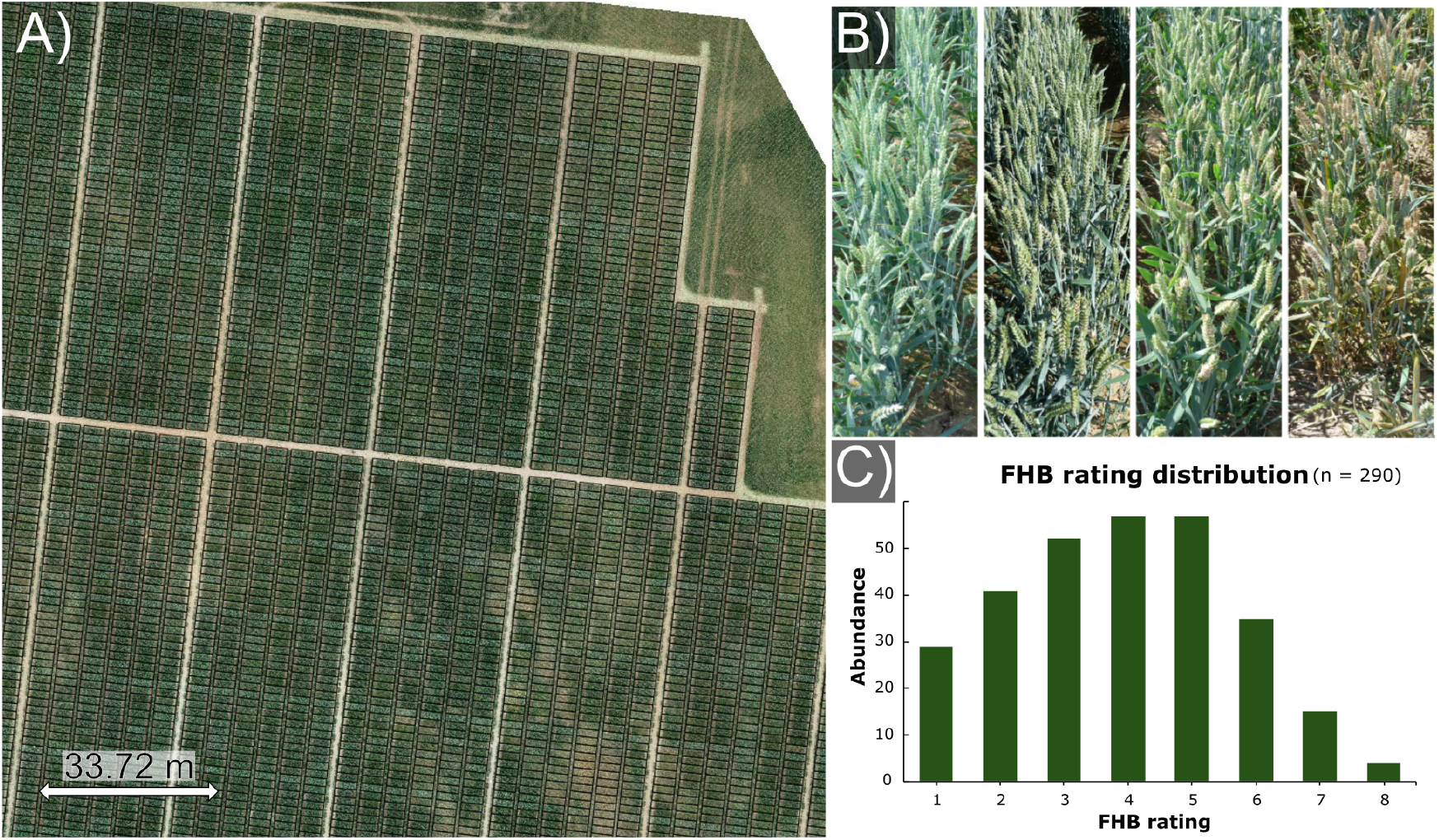
Drone image overview of the field trial with marked plots (A), FHB severities in increasing order (B), and distribution of FHB ratings for the 145 plots analysed in duplicate (C). Panel (A) is published with explicit permission and under the copyright of geo-konzept GmbH, Adelschlag, Germany. Panel (B) is adapted from Rößle *et al*. (10) and used under the terms of the CC BY 4.0 licence.

A detailed summary of *Fusarium* infection and visual disease progression is provided in **Supplementary Table S1**.

### Mycotoxin quantification

The targeted LC–MS/MS screening of 23 mycotoxins revealed an expected dominance of *Fusarium*-derived mycotoxins. Among these, DON and its related metabolites, 3-AcDON and DON-3G, showed the highest concentrations, reflecting their central roles as the primary toxin of *F. culmorum*, its biosynthetic precursor, and the plant’s glucosylation-based detoxification product. Other *Fusarium* toxins, including enniatins (A, A1, B, B1), beauvericin, and zearalenone, occurred only sporadically and at low abundance, consistent with background co-infection by other *Fusarium* species, while nivalenol and fusarenone X were absent. *Alternaria* contamination appeared unstructured across samples, with TeA consistently detected at the highest levels, serving as the principal marker of *Alternaria* activity in the field. The quantification results for DON, 3-AcDON, DON-3G, and TeA, as well as the respective infestation severity ratings are given in **Supplementary Table S1**. DON was detected in all samples and spanned a wide range of 2–5,000 µg kg^-1^, reflecting the naturally occurring variability in cultivar susceptibility and the expected heterogeneity of infection progress under field conditions.

The observed DON concentrations fall within the range reported across multiple wheat field studies—both inoculated and naturally infected—indicating that the spectrum captured in our trial is well aligned with conditions encountered in agronomic practice (17, 18, 51). The conjugated detoxification product DON-3G typically represented 5–20% of the corresponding DON content, whereas 3-AcDON occurred at lower levels. Although the DON-3G-to-DON ratio varied among samples, the glycoside’s relative abundance remained too low to indicate a major contribution to resistance. Deliberate inoculation led to DON concentrations approaching 5,000 µg kg^-1^, a toxin burden far above food and feed safety limits. According to Regulation (EU) 2024/1022, the maximum permitted DON concentration in unprocessed cereals is 1,000 µg kg^-1^ (6). In this study, 41% of samples exceeded this limit, increasing to 46% when considering the modified forms DON-3G and 3-AcDON (both expressed as DON equivalents; no legal thresholds).

Natural *Alternaria* infestation was monitored across all samples, with TeA emerging as the predominant metabolite, detected in 140 of 145 samples at concentrations ranging from 0.79 to 60 µg kg^-1^. Its abundance showed no correlation with *Fusarium*-derived toxins or overall grain quality. Tentoxin was found in about two-thirds of the samples, though concentrations remained largely below the LOQ of 0.46 µg kg^-1^, with a maximum concentration detected of 1.2 µg kg^-1^. Other *Alternaria* metabolites, including alternariol and alternariol monomethyl ether, were absent. The generally low contamination levels likely reflect the dry 2022 growing season, limiting conclusions about potential competitive or sequential fungal interactions.

### Relationship between visual FHB score and toxin concentration

To assess the relationship between visually determined FHB severity as the proportion of visibly infected ears, and toxin concentration as an indicator of infestation progression, post-hoc pairwise comparisons were performed using the Games–Howell test after Welch’s ANOVA and summarized into Compact Letter Displays **(Supplementary Table S3)**. DON concentrations increased significantly from FHB classes 1 to 2 and 3 to 4 (Games–Howell, p < 0.05). Beginning at class ≥4, however, DON values entered a broad high-level plateau: severity classes 4–8 did not differ significantly from each other, indicating that visual scores ≥ 4 no longer provide quantitative resolution of final toxin burden. Accordingly, Compact Letter Display grouping separated low-severity classes (1–3) from uniformly high DON levels observed at severity ≥4. Respective toxin contents (µg kg^-1^) are plotted against the visual quality ratings in **Figure 2**. To account for the broad dynamic range of toxin concentrations, a logarithmic scale was applied for visualization purposes only. Beyond the threshold of FHB rating 3 (5% of infected spikelet), visual assessment no longer provided a reliable indication of the toxin burden of the ripe grain relevant for food safety and marketability. In resistance studies, low FHB severity scores are often associated with reduced DON accumulation in mature grain and interpreted as indicators of effective early containment (52–54). However, a consistent quantitative relationship between visual severity and the wide range of DON concentrations observed under field conditions has not been reliably demonstrated (53–57), and our field data similarly revealed no such consistent association. The *Alternaria* toxin TeA, found due to natural infestation, displayed no systematic relationship with neither FHB severity nor *Fusarium* toxin load.

Visual FHB assessments were performed during milk ripeness, when symptoms are most reliably distinguished in standard field practice. Grain for toxin and metabolomic analysis was collected later at full physiological maturity, resulting in a temporal offset between symptom scoring and biochemical profiling. Consequently, the visual ratings capture an early and phenology-dependent snapshot of symptom expression, resulting in heterogeneous assessments that offer limited insight into the final toxin burden and the plant’s metabolic status at harvest. To capture biologically meaningful relationships, further correlations with the metabolome were based on measured toxin concentrations rather than visual FHB scores. Mycotoxin concentrations capture the cumulative progression of fungal activity and toxin-associated stress and therefore offer a temporally relevant and quantitative biochemical measure of pathogen impact.

**Figure 2.**
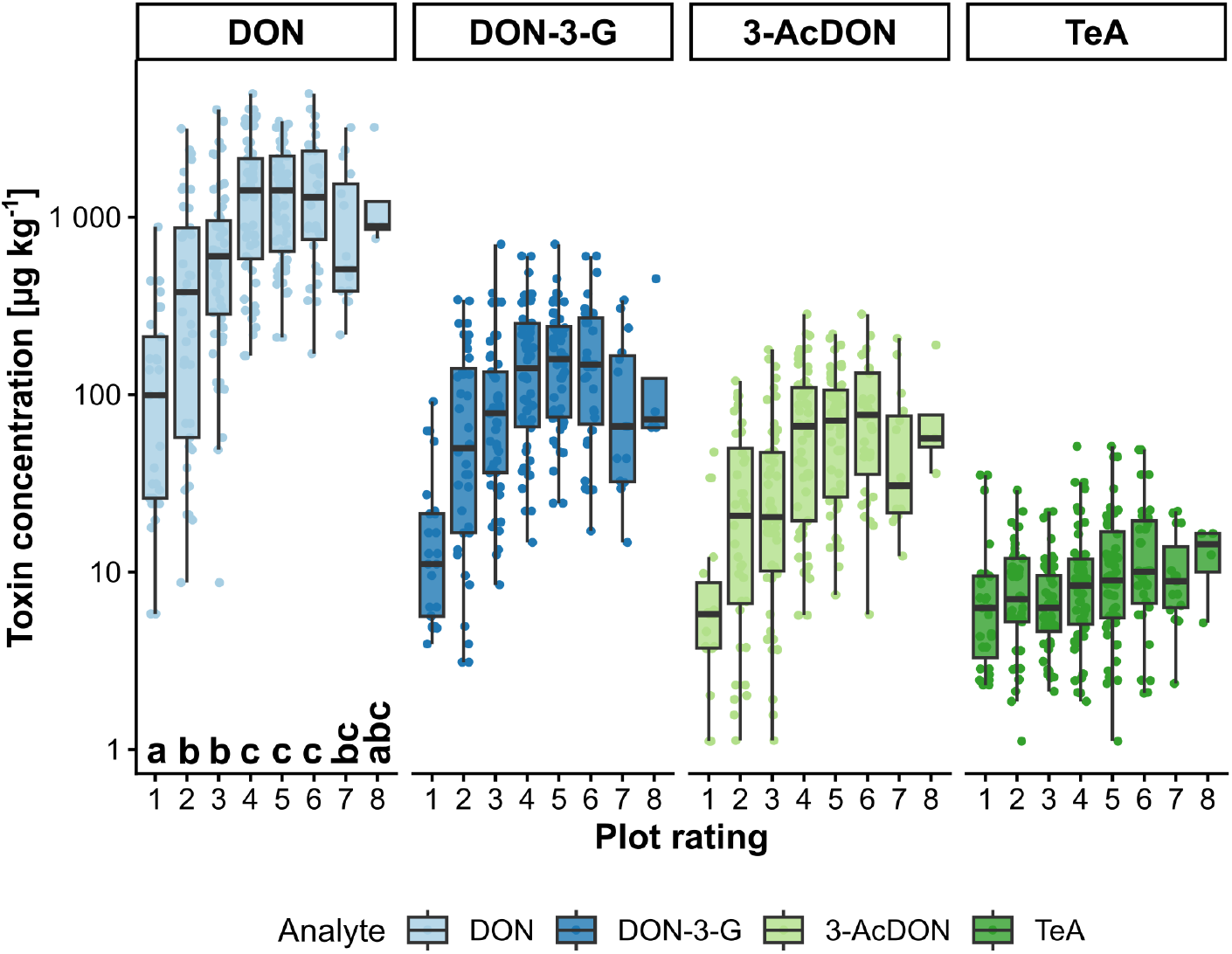
Mycotoxin concentration (y-axis, log-scale) against the visual FHB rating (x-axis) for DON, DON-3-G, 3-Ac-DON, and TeA. Severity scores 1–3 reflect lower and increasing DON levels, whereas severity *≥*4 shows uniformly high DON without significant separation among classes, as indicated by Compact Letter Display (bottom left). Statistics are detailed in **Supplementary Table S3**. Values below LOQ are not shown.

### Metabolomics data and statistics

QuEChERS extracts were analyzed in duplicates using reversed-phase UPLC-ToF mass spectrometry (**Supplementary Figure S1A**). After data processing, an average of 2,905 *±* 335 chromatographic features were obtained for each extract, resulting in a dataset spanning 6,829 features. The previously quantified mycotoxins remained below the LOD of the non-targeted system. Both coumaroylagmatine and coumaroylputrescine, reported to be linked to the resistance gene *TaACT* (25), were not detected in our experimental setup. The overall variability within the dataset was assessed using PCA, which showed no distinct correlation between toxin levels and metabolomic profiles in the unsupervised statistical analysis (**Supplementary Figure S1B**). Considering the diversity of wheat genotypes examined and their unique genetic backgrounds, a high degree of variability in metabolic patterns was expected. The low explained variance in the first two principal components (PC1: 12.7%, PC2: 6.1%) reflects the high metabolic heterogeneity within the field experiment, driven by the diverse genetic backgrounds and nutritional microenvironments of the wheat genotypes. This complexity captures the biological reality of field conditions and allows the identification of metabolic responses that persist across genotypes and environmental variation.

To disentangle these universal responses from background variability, we applied a supervised OPLS model using the measured DON concentration as the dependent y-variable. The OPLS model showed strong predictive performance, with an R^2^Y of 0.98 and a Q^2^ of 0.95, indicating excellent model fit and predictive reliability. Overfitting was excluded by CV-ANOVA (p ≪ 0.05), confirming the robustness of the relationship between metabolite profiles and DON concentrations (46, 51) (**Supplementary Table S8**). The corresponding score plot (**Figure 3A**) shows a clear gradient of mycotoxin load along the first predictive component, indicating that the model effectively captures metabolic processes linked to the progression and biochemical impact of *F. culmorum* infection. These responses occur consistently across the diverse genotypes studied, reflecting shared physiological mechanisms rather than cultivar-specific effects. Unlike the ambiguous relationship between visual disease symptoms and toxin exposure, measured toxin load provides a robust quantitative indicator of the metabolic processes associated with fungal activity and host response.

**Figure 3.**
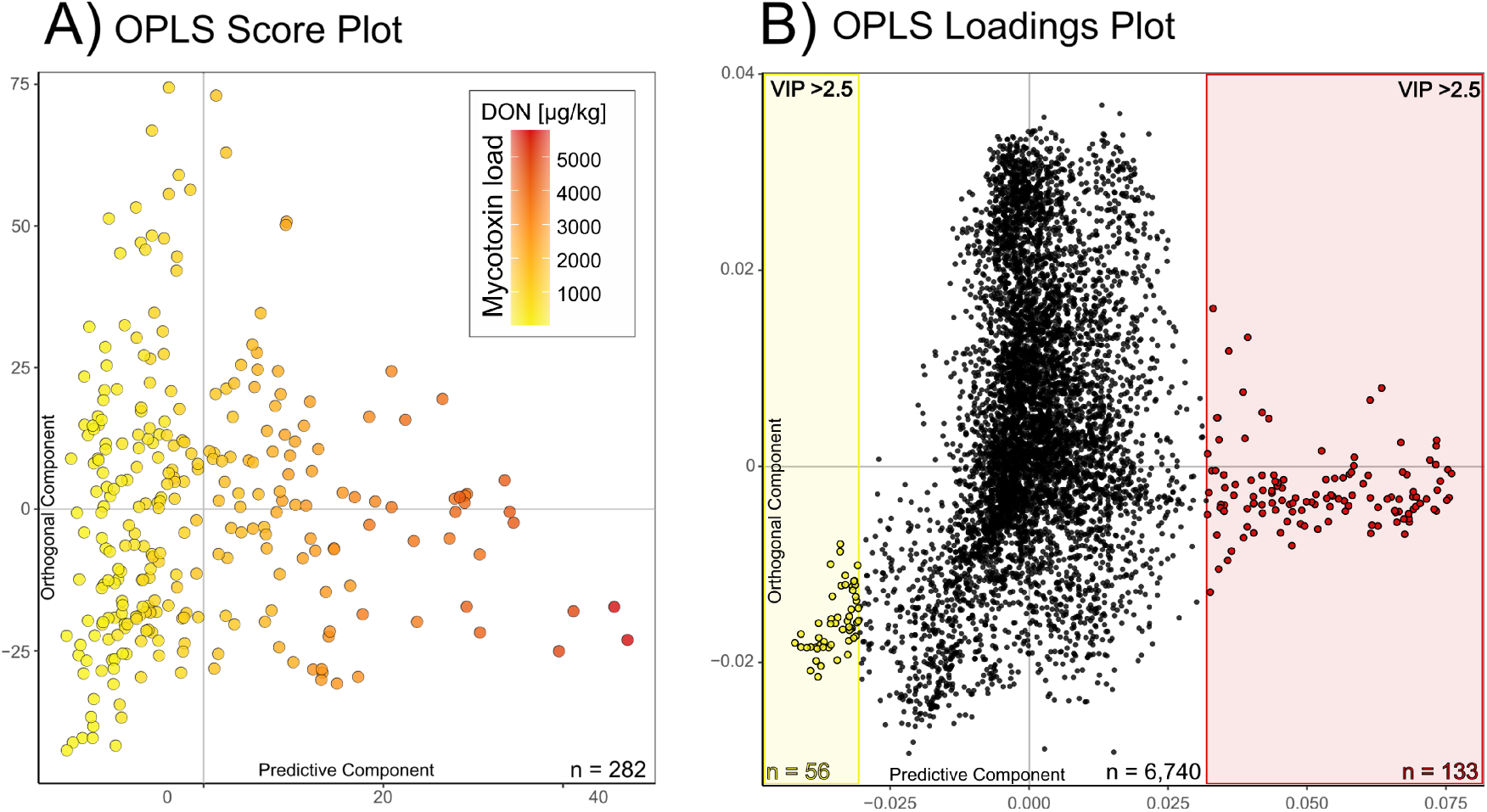
Score (A) and loadings (B) plot of the OPLS model of the metabolomics data utilizing the DON-concentration as the dependant y-variable. Each sample replicate is represented as a dot, coloured according to the measured DON concentration, ranging from 10 µg kg^-1^ (yellow) to 4,960 µg kg^-1^ (red) (A). Each chromatographic feature is represented as a dot, with those most significantly correlated with the y-variable (VIP *>* 2.5) highlighted in yellow (negative) and red (positive) (B).

The strong correlation between metabolomic patterns and measured toxin loads, despite the toxins themselves not being directly detected in the untargeted dataset, demonstrates that the overall metabolic signature reliably reflects toxin accumulation. Positively correlated features showed linear relationships with DON concentration, confirmed by correlation and regression analyses (*α* = 0.05) (**Supplementary Figure S2A**). Regression modelling using linear, LASSO, and PLS approaches, based on a data split into training (70%), test (20%), and validation (10%) subsets, yielded highly accurate predictions of DON concentrations from metabolomic features, achieving coefficients of determination (R^2^) of up to 0.98 (**Supplementary Figure S3**). The loadings plot (**Figure 3B**) highlights the metabolites most strongly associated with toxin accumulation, selected using a stringent and statistically robust VIP threshold of 2.5. This higher cutoff, well above the conventional VIP > 1 used in many metabolomics studies, was chosen to isolate only the most influential features and thereby enable biologically meaningful interpretation. A substantial number of highly significant metabolites (VIP > 6) showed positive correlation with DON levels (positive loadings), suggesting a fungal origin and reflecting processes linked to successful infection. In contrast, negatively correlated features (negative loadings) likely represent host-derived defense or interaction metabolites. The structural characterization and biological interpretation of these key compounds are described in the following section.

### Metabolic pathways linked to infection severity

To move from statistical association to biochemical interpretation, we resolved the compound class of metabolites linked to toxin accumulation. Tandem mass spectral data exported from MS-DIAL were analyzed in the SIRIUS (45) environment, where fragmentation-based molecular formula prediction, compound class annotation, and structural inference enabled the contextualization of these features within metabolite pathways. Structural and chemical classification of metabolites was performed using CANOPUS (Class Assignment and Ontology Prediction Using Mass Spectrometry), a deep-learning framework trained on nearly 25,000 reference compounds from NIST 2017, GNPS, and MassBank (48). The twenty most significant features showing positive or negative correlation with toxin levels are summarized in **Table 1**, and the corresponding MS^2^ spectra are provided in the Supplementary Information (**Supplementary Tables S9 and S10**).

**Table 1.**
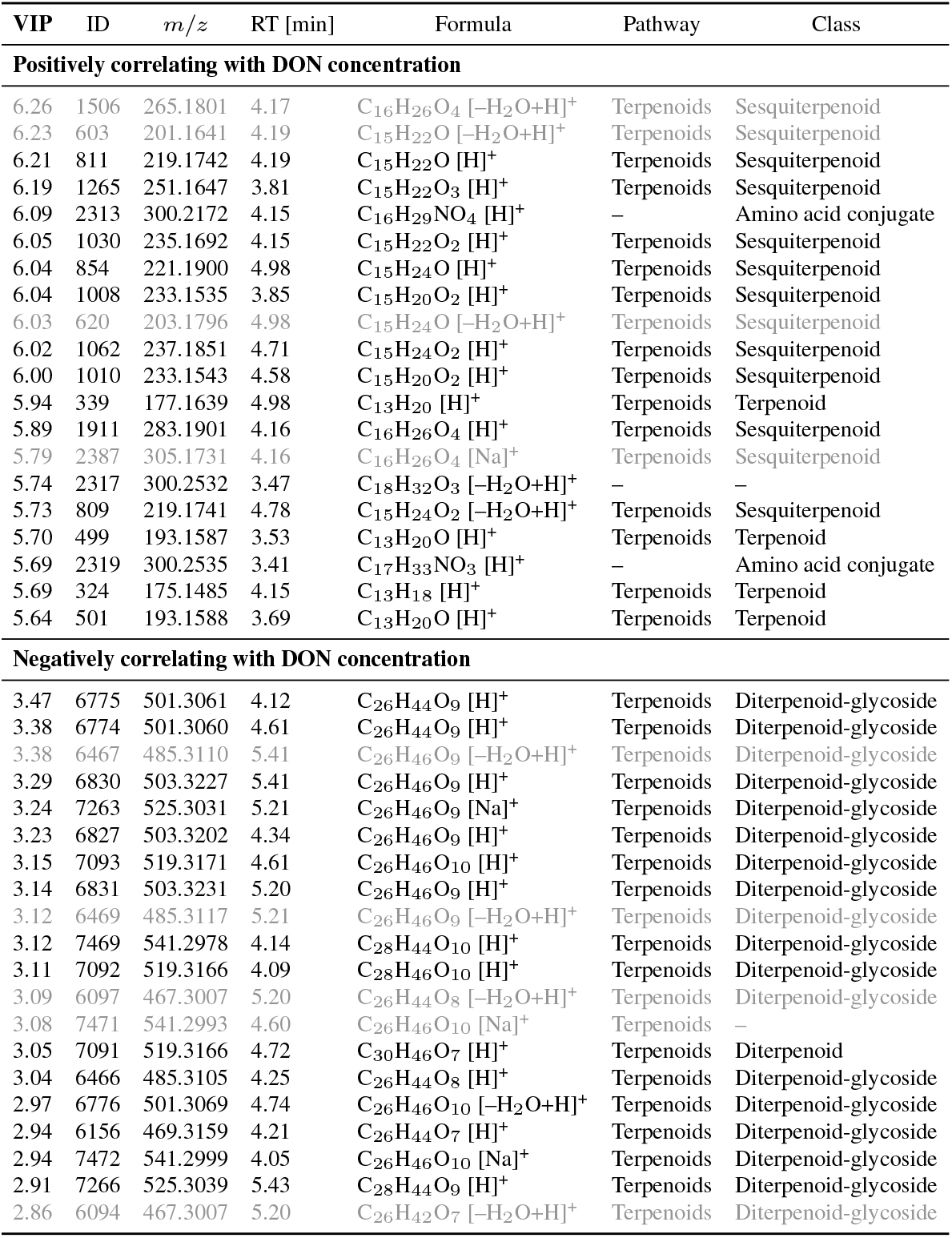
The most significant features positively (top) and negatively (bottom) correlated with the DON concentration quantified in the wheat grain samples. The VIP values of the OPLS model, m/z values, retention time, molecular formula and ionization, annotated pathway (Sirius ClassyFire), and compound class (Sirius CANOPUS). Ambiguous classes are not reported. Compounds appearing as duplicates due to ion-source artefact and adduct formation (-H_2_ O and +Na^+^) are shown in grey.

The metabolites that increased most strongly with the progression of *F. culmorum* infection were consistently linked to sesquiterpene metabolism. The sesquiterpenoid annotation in the CANOPUS framework is based on their characteristic C_15_ backbone and diagnostic MS^2^ fragment ions (e.g., *m/z* 93.0700 [C_7_H_9_]^+^, *m/z* 105.0700 [C_8_H_9_]^+^, *m/z* 109.1012 [C_8_H_12_]^+^, or *m/z* 121.1012 [C_9_H_12_]^+^) (identification level 3) (50). Sesquiterpenes are farnesyl-derived C_15_-terpenoids central to *Fusarium* virulence, with trichothecenes constituting their most potent phytotoxic representatives. Among these, DON and its derivatives—our dependent variable that is not part of the untargeted metabolomics dataset—are well established as key virulence factors that disrupt host protein synthesis and critically determine disease severity (58–60). Several sesquiterpene intermediates associated with trichothecene biosynthesis increased in abundance with higher toxin load, including hydroxytrichodiene (C_15_H_24_O, level 2a), isotrichodiol (C_15_H_22_O_2_, level 2b), isotrichotriol/trichodermol (C_15_H_22_O_3_, level 2b), isotrichodermin (C_15_H_22_O_4_, level 3), decalonectrin / calonectrin (C_15_H_22_O_5_, level 2b), and nortrichodienes (C_13_H_18_, C_13_H_20_, C_13_H_20_O, level 3). These annotations were supported by *in silico* fragmentation analysis via CSI:FingerID (49) and verified against reference spectra in the MassBank of North America (University of California, Davis). The pronounced enrichment of sesquiterpene-derived metabolites mirrors established *Fusarium* infection pathways, and their near-linear relationship with DON concentrations reinforces the mechanistic link between metabolic signatures and pathogen virulence.

In contrast, the biochemical origin and functional role of metabolites associated with constitutive or inducible plant defence processes are more difficult to define. These compounds, which were consistently enriched at low toxin concentrations, predominantly mapped to diterpene metabolism (**Table 1**). Their shared C_20_ backbone, diagnostic fragment ions indicative of an extended terpenoid core (e.g., *m/z* 321.2424 [C_20_H_33_O_3_]^+^, *m/z* 305.2480 [C_20_H_33_O_2_]^+^, *m/z* 209.1900 [C_14_H_24_O]^+^, or *m/z* 149.1325 [C_11_H_17_]^+^), and recurring hexosyl neutral loss (*−*162.0534 [C_6_H_10_O_5_]) classify them as glycosylated diterpenes (level 3). These metabolites showed their highest intensities in samples with minimal DON contamination, and their inverse association diminished progressively as toxin load increased (**Supplementary Figure S2B**). Since their abundance decreases with rising fungal activity, contrasting the accumulation of fungal metabolites such as DON and DON-3-G, a fungal origin with subsequent plant-mediated glycosylation is improbable, though not formally excluded. The consistent detection of these glycosylated metabolites, together with their association with effective pathogen containment, supports their origin as endogenous components of wheat secondary metabolism.

Previous work by Gauthier *et al*. (61) summarized metabolic signatures associated with cereal defence against *Fusarium*, including diterpenoid accumulation in barley (62–64). These metabolites, however, represented only a minor fraction of the reported defence-associated profiles and were characterized mainly under controlled laboratory conditions, leaving their ecological relevance unresolved. In a recent comprehensive transcriptomic analysis across a broad panel of wheat genotypes, Buerstmayr *et al*. (65) reported elevated expression of terpene-related pathways associated with resistance and early containment of fungal colonization. In Maize and rice, diterpenoids are well established as phytoalexins, where they disrupt fungal membranes following chitin recognition (66, 67). The underlying labdane-related diterpenoid metabolism is thought to have diversified early in plant evolution (68), giving rise to distinct ent-kaurene synthase-like (KSL) gene families (69). Recent genetic and transcriptomic studies have extended this concept to the *Pooideae*, suggesting that diterpene-mediated defense mechanisms may also operate in wheat and barley. Polturak *et al*. (70) identified two biosynthetic gene clusters in *Triticum aestivum* co-expressed with terpene synthases, likely sharing an evolutionary origin with the momilactone phytoalexin cluster of rice. Our findings provide complementary evidence from non-targeted metabolomics under field conditions, supporting the hypothesis that the diterpene metabolism of wheat plays a functional role in defence against *Fusarium* infection.

A consistent association between glycosylated diterpenes and reduced effective pathogen activity was observed across genotypes. In cereals, diterpenoids are known precursors of diverse antimicrobial phytoalexins; however their occurrence mainly as glycosides suggests additional layers of regulation. Such glycosylation may serve multiple, not mutually exclusive functions, including modulation of bioactivity, intracellular storage, and spatial or temporal control of diterpenoid availability. While it remains unclear whether these compounds function as phytoalexins, phytoanticipins, or represent inert storage forms, their strong association with early containment of infection severity points to a close link with effective host–pathogen interaction outcomes. In other plant systems, glycosylation has been proposed to either enable stress-inducible hydrolysis of conjugates to release bioactive aglycones or to confer biological activity directly to the glycosylated metabolites by improving solubility, stability, or transport properties (71). The accumulation of diterpene glycosides, particularly in samples with low *Fusarium* infection efficiency, may indicate an effective host response in which (glycosylated) diterpenes form part of an early containment strategy. This pattern points to a potential biochemical role of diterpene metabolism in plant defence, warranting further investigation into the biosynthetic origin, enzymatic mediators, and functional significance of diterpenes and their glycosides in the wheat–*Fusarium* interaction.

## Conclusion

Our field study integrated multiple *Triticum aestivum* genotypes grown under agronomically realistic conditions and deliberately inoculated with *Fusarium culmorum*, generating a broad and biologically relevant gradient of infection severities and toxin loads that reflected natural field variability. By combining accurate mycotoxin quantification with untargeted metabolomics across genetically diverse wheat material, we identified infection-associated metabolic signatures that remained consistent across genotypic and environmental heterogeneity. Using mycotoxin concentrations as quantitative, continuous measures overcame key limitations of conventional visual disease scoring and enabled robust associations between metabolic processes and infection intensity. The strong co-variation of sesquiterpene-derived metabolites with toxin accumulation underscores their close linkage to fungal virulence pathways and warrants further investigation of trichothecene precursor dynamics and signalling processes. Conversely, the conserved elevated presence of glycosylated diterpene conjugates at low toxin loads points to a constitutive or early-acting metabolic containment mechanism intrinsic to the wheat host that had so far remained largely unexplored. Elucidating the biochemical function, biosynthetic enzymes, and regulatory networks underlying these diterpene conjugates represents a critical next step toward understanding potential diterpene-mediated defence strategies in wheat. Collectively, our findings demonstrate that integrating metabolomic profiling with quantitative toxin measurements provides a powerful framework for disentangling complex plant–pathogen interactions under field conditions.

## DATA AVAILABILITY

The mycotoxin quantification raw data will be made available upon reasonable request, without undue reservation. The LC-ToF-MS data set and mgf file were uploaded to the MassIVE (UC San Diego) repository (http://doi.org/10.25345/C52B8VR38) and the mandatory password is made available upon request.

## AUTHOR CONTRIBUTIONS

Software: S.A.P., J.S.S., P.S.K.; Formal analysis: S.A.P., F.D., M.B., L.R., E.K.; Investigation: S.A.P., F.D., M.B., L.R.; Resources: A.K., L.P., J.S.S., P.O.N., P.S.K., M.R.; Visualization: S.A.P.; F.D.; M.B., J.S.S.; Supervision: S.A.P., F.D., P.O.N., S.A., P.S.K., M.R.; Project administration: S.A.P., F.D., A.H., J.S.S., P.O.N., M.R.; Funding acquisition: S.A.P., J.S.S., A.H., P.O.N., M.R., Writing – Original Draft: S.A.P., F.D.; Conceptualization, Methodology, Validation, Writing – Review, Editing & Approval: all authors.

