## Supplementary Information for "Glycosylated diterpenes associate with early containment of *Fusarium culmorum* infection across wheat (*Triticum aestivum* L.) genotypes under field conditions"

#### Contents

**Supplementary Table S1.** Overview of the field study, including parcel number (BEO), wheat variety/genotype, FHB scoring (FHB1 and FHB2), and toxin concentrations (n.d. = not detected, < LOQ = below quantification limit)........ 4

**Supplementary Table S2.** LC-MS/MS parameters of the analyzed mycotoxins. Detailed are the analyte, its ionization state, precursor mass-to-charge-ratio, two transition product ions mass-to-charge-ratios, respective voltage focusing before Q1, voltage in the collision cell, voltage applied before Q3, and the retention time. Abbreviations in chronological order: AOH = alternariol, AME = alternariol monomethylether, -G = respective glucosides, -S = respective sulfates, ATX I = altertoxin I, TeA = tenuazonic acid, TEN = tentoxin, DON = deoxynivalenol, 3-AcDON = 3-acetyl-DON, NIV = nivalenol, Fus X = fusarenone X, T-2 = T-2 toxin, HT-2 = HT-2 toxin; ZEN = zearalenone, ENN = enniatin, BEA = beauvericin. ........ 9

#### Supplementary Tables

**Supplementary Table S1.** Overview of the field study, including parcel number (BEO), wheat variety/genotype, FHB scoring (FHB1 and FHB2), and toxin concentrations (n.d. = not detected, < LOQ = below quantification limit).

| BEO | Variety / Genotype | FHB 1 | FHB 2 | DON [µg kg$^{-1}$] | DON-3-G [µg kg$^{-1}$] | 3-AcDON [µg kg$^{-1}$] | TeA [µg kg$^{-1}$] |
|---|---|---|---|---|---|---|---|
| **1** | JB Asano | 5 | 5 | 732.45 | 73.17 | 25.11 | 12.56 |
| **2** | Dichter | 1 | 2 | 19.58 | n.d. | n.d. | 7.02 |
| **3** | 13360r4 | 3 | 3 | 116.68 | 19.10 | 3.64 | 3.91 |
| **4** | 13360r6 | 3 | 3 | 626.79 | 84.38 | 20.85 | 5.00 |
| **5** | 13360r8 | 2 | 2 | 68.56 | 16.58 | 3.75 | 2.85 |
| **6** | 13360r9 | 2 | 2 | 30.33 | n.d. | <LOQ | 2.47 |
| **7** | 13360s1 | 4 | 3 | 343.98 | 47.84 | 16.31 | 4.05 |
| **8** | 13593r5 | 6 | 6 | 2362.79 | 228.61 | 132.49 | 17.40 |
| **9** | 13593r7 | 2 | 3 | 8.69 | n.d. | 1.56 | 10.00 |
| **10** | 13602r2 | 3 | 3 | 107.02 | n.d. | 4.17 | 6.55 |
| **11** | 13602r3 | 2 | 3 | 57.15 | 12.48 | 1.91 | 7.27 |
| **12** | 13602r4 | 3 | 4 | 267.00 | 29.50 | 9.15 | 5.15 |
| **13** | 13629r9 | 3 | 3 | 289.20 | 36.28 | 10.22 | 6.26 |
| **14** | 13732t9 | 2 | 2 | 777.96 | 129.48 | 27.66 | 5.49 |
| **15** | 13834r9 | 3 | 2 | 544.13 | 98.32 | 18.31 | 2.82 |
| **16** | 13854s3 | 4 | 4 | 297.62 | 34.96 | 12.04 | 9.17 |
| **17** | 13854s4 | 3 | 3 | 623.82 | 78.29 | 19.88 | 5.63 |
| **18** | 13854s5 | 2 | 3 | 219.38 | 30.86 | 9.27 | 11.88 |
| **19** | 13854s6 | 2 | 3 | 1143.76 | 134.28 | 46.60 | 12.19 |
| **20** | 14381p1 | 6 | 5 | 417.89 | 81.52 | 26.41 | 18.61 |
| **21** | 14381p2 | 5 | 6 | 811.25 | 171.66 | 79.20 | 3.89 |
| **22** | 14398p3 | 3 | 5 | 3460.18 | 703.02 | 143.67 | 2.55 |
| **23** | 14398p7 | 3 | 4 | 899.00 | 164.56 | 34.63 | 2.67 |
| **24** | 14398p8 | 2 | 3 | 824.28 | 199.75 | 43.24 | 2.12 |
| **25** | 14411p4 | 2 | 3 | 419.48 | 30.06 | 10.69 | 9.42 |
| **26** | 14411p5 | 2 | 3 | 291.25 | 17.92 | 9.75 | 10.71 |
| **27** | JB Asano | 4 | 5 | 2676.50 | 156.05 | 103.10 | 23.07 |
| **28** | Dichter | 2 | 2 | 21.02 | <LOQ | 2.30 | 17.96 |
| **29** | 14411r7 | 4 | 3 | 579.02 | 50.79 | 16.67 | 10.78 |
| **30** | Dichter | 1 | 2 | 38.49 | <LOQ | 0.81 | 28.80 |
| **31** | 14414p3 | 4 | 4 | 290.83 | 38.42 | 9.91 | 12.51 |
| **32** | 14418p4 | 3 | 5 | 1263.73 | 123.65 | 37.65 | 4.86 |
| **33** | JB Asano | 5 | 6 | 1900.63 | 147.03 | 84.90 | 20.17 |
| **34** | 14420r4 | 4 | 4 | 798.46 | 81.17 | 35.80 | 10.25 |
| **35** | 14421p3 | 2 | 4 | 792.73 | 64.88 | 23.29 | 5.81 |
| **36** | 14434p2 | 3 | 5 | 2274.88 | 331.33 | 93.53 | 3.13 |
| **37** | 14434p4 | 4 | 6 | 690.38 | 73.88 | 28.24 | 2.07 |

**Supplementary Table S1.** Continued.

| BEO | Variety | FHB 1 | FHB 2 | DON [µg kg$^{-1}$] | DON-3-G [µg kg$^{-1}$] | 3-AcDON [µg kg$^{-1}$] | TeA [µg kg$^{-1}$] |
|---|---|---|---|---|---|---|---|
| **38** | 14434p5 | 3 | 4 | 531.39 | 69.05 | 19.39 | 3.92 |
| **39** | 14434r1 | 5 | 5 | 372.56 | 46.93 | 13.69 | 3.87 |
| **40** | 14434r2 | 5 | 5 | 209.53 | 24.40 | 7.42 | 8.56 |
| **41** | Dichter | 1 | 2 | 28.87 | n.d. | <LOQ | 14.37 |
| **42** | JB Asano | 4 | 4 | 1300.18 | 122.52 | 42.83 | 31.95 |
| **43** | Dichter | 1 | 1 | 24.18 | n.d. | <LOQ | 6.25 |
| **44** | 14448p6 | 5 | 4 | 424.51 | 36.90 | 10.69 | 6.13 |
| **45** | Floki | 5 | 4 | 498.93 | 71.79 | 15.46 | 9.37 |
| **46** | 14448p9 | 5 | 5 | 573.69 | 74.95 | 28.25 | 44.49 |
| **47** | 14451p4 | 4 | 4 | 165.58 | 22.02 | 5.69 | 8.37 |
| **48** | JB Asano | 4 | 5 | 2631.63 | 217.84 | 135.03 | 22.14 |
| **49** | 14501p2 | 1 | 1 | 159.18 | 16.63 | 5.78 | 7.13 |
| **50** | 14501p9 | 1 | 1 | 26.04 | <LOQ | <LOQ | 0.97 |
| **51** | 14501r7 | 2 | 4 | 2279.28 | 251.14 | 119.33 | 12.74 |
| **52** | 14513p1 | 3 | 5 | 983.07 | 69.73 | 48.30 | 11.90 |
| **53** | 14513p6 | 2 | 4 | 1472.46 | 125.71 | 80.66 | 9.91 |
| **54** | 14513r1 | 3 | 5 | 519.80 | 63.33 | 22.61 | 5.05 |
| **55** | 14513r2 | 2 | 5 | 377.63 | 47.42 | 14.38 | 13.55 |
| **56** | 14529p1 | 4 | 5 | 1649.68 | 190.86 | 66.30 | 12.01 |
| **57** | 14529p3 | 3 | 5 | 639.39 | 82.86 | 15.08 | 10.78 |
| **58** | JB Asano | 3 | 5 | 2249.57 | 215.04 | 108.75 | 21.73 |
| **59** | 14544p1 | 3 | 4 | 4022.01 | 329.31 | 158.76 | 4.83 |
| **60** | 14551p2 | 1 | 1 | 137.85 | 21.23 | 3.69 | 2.30 |
| **61** | 14551p4 | 1 | 1 | 99.03 | 12.67 | 3.79 | 2.45 |
| **62** | 14551p5 | 1 | 1 | 5.80 | <LOQ | <LOQ | 4.33 |
| **63** | 14551r1 | 1 | 1 | 31.30 | <LOQ | 1.11 | 9.77 |
| **64** | 14553p1 | 1 | 1 | 311.11 | 11.04 | 5.87 | 3.77 |
| **65** | 14626p6 | 3 | 5 | 655.84 | 128.28 | 31.13 | 6.32 |
| **66** | 14639p4 | 2 | 4 | 2104.36 | 341.98 | 87.41 | 1.86 |
| **67** | JB Asano | 5 | 4 | 1500.66 | 183.47 | 67.68 | 16.33 |
| **68** | JB Asano | 4 | 5 | 1911.88 | 266.17 | 87.37 | 22.06 |
| **69** | 14389p6 | 4 | 4 | 3578.49 | 362.44 | 179.63 | 4.33 |
| **70** | JB Asano | 2 | 4 | 2391.16 | 336.31 | 101.44 | 18.99 |
| **71** | 14389p5 | 4 | 6 | 4960.47 | 602.69 | 214.46 | 9.18 |
| **72** | 14420r8 | 1 | 1 | 17.76 | <LOQ | <LOQ | 35.05 |

**Supplementary Table S1.** continued.

| BEO | Variety | FHB 1 | FHB 2 | DON [µg kg$^{-1}$] | DON-3-G [µg kg$^{-1}$] | 3-AcDON [µg kg$^{-1}$] | TeA [µg kg$^{-1}$] |
|---|---|---|---|---|---|---|---|
| **73** | JB Asano | 4 | 5 | 3273.64 | 332.18 | 147.99 | 11.72 |
| **74** | Informer | 1 | 2 | 247.56 | 22.09 | 12.11 | 9.52 |
| **75** | RGT Reform | 1 | 3 | 210.97 | 27.20 | 4.61 | 8.58 |
| **76** | LG Initial | 2 | 3 | 466.52 | 40.36 | 25.02 | 21.90 |
| **77** | KWS Emerick | 1 | 3 | 378.77 | 54.35 | 9.81 | 2.64 |
| **78** | Campesino | 5 | 6 | 1740.57 | 285.72 | 74.71 | 2.45 |
| **79** | Floki | 3 | 4 | 1511.97 | 166.13 | 68.05 | 9.16 |
| **80** | KWS Donovan | 2 | 3 | 870.25 | 83.04 | 35.17 | 5.43 |
| **81** | Ellvis | 2 | 3 | 48.76 | 8.46 | 1.12 | 7.31 |
| **82** | JB Diego | 2 | 4 | 1428.40 | 180.74 | 60.14 | 5.37 |
| **83** | JB Asano | 5 | 4 | 3252.50 | 332.30 | 143.22 | 18.33 |
| **84** | Trapez | 4 | 6 | 3716.58 | 486.42 | 181.16 | 6.80 |
| **85** | Inspiration | 1 | 4 | 880.38 | 91.11 | 47.37 | 4.49 |
| **86** | Dichter | 1 | 2 | 51.00 | <LOQ | 2.01 | 5.74 |
| **87** | Brentano | 2 | 3 | 1432.75 | 216.42 | 61.49 | 5.22 |
| **88** | Bosporus | 1 | 1 | 436.68 | 62.27 | 33.86 | 2.84 |
| **89** | Maradona | 1 | 2 | 155.90 | 9.53 | 6.09 | 9.44 |
| **90** | Terence | 2 | 3 | 471.79 | 52.44 | 23.92 | 10.35 |
| **91** | Akzent | 3 | 2 | 198.16 | 13.32 | 10.91 | 15.36 |
| **92** | Asgard | 2 | 2 | 147.19 | 35.28 | 6.81 | 3.60 |
| **93** | Damian | 4 | 6 | 4058.43 | 600.41 | 283.59 | 2.09 |
| **94** | Shaun | 4 | 5 | 1628.73 | 252.76 | 109.02 | 9.05 |
| **95** | 11713a33 | 3 | 5 | 2457.73 | 331.86 | 123.75 | 2.77 |
| **96** | 12286p61 | 4 | 5 | 1409.88 | n.d. | 78.91 | n.d. |
| **97** | 12337a6 | 3 | 4 | 710.01 | 93.74 | 41.39 | 6.79 |
| **98** | 13355p21 | 4 | 5 | 3344.24 | 368.29 | 219.24 | 8.88 |
| **99** | Floki | 2 | 4 | 766.56 | 100.68 | 41.55 | 6.24 |
| **100** | 13079p223 | 4 | 6 | 865.21 | 125.87 | 66.11 | 2.44 |
| **101** | 13217p412 | 2 | 4 | 1795.19 | 217.54 | 97.80 | 6.07 |
| **102** | JB Asano | 4 | 5 | 2076.48 | 223.84 | 116.40 | 50.94 |
| **103** | 13313r313 | 3 | 4 | 314.76 | 42.27 | 17.74 | 7.65 |
| **104** | 12411a441 | 2 | 5 | 1121.28 | 160.46 | 69.19 | 5.61 |
| **105** | 12569c143 | 3 | 5 | 1032.84 | 162.04 | 63.07 | 5.58 |
| **106** | 12687c743 | 4 | 5 | 1629.96 | 235.22 | 113.24 | 7.52 |

**Supplementary Table S1.** continued.

| BEO | Variety | FHB 1 | FHB 2 | DON [µg kg$^{-1}$] | DON-3-G [µg kg$^{-1}$] | 3-AcDON [µg kg$^{-1}$] | TeA [µg kg$^{-1}$] |
|---|---|---|---|---|---|---|---|
| **107** | 12696b115 | 3 | 4 | 238.20 | 38.20 | 16.73 | 3.66 |
| **108** | Floki | 3 | 5 | 2654.21 | 371.88 | 178.94 | 9.53 |
| **109** | 13648r63 | 2 | 5 | 3143.03 | 251.68 | 96.04 | 1.11 |
| **110** | 12865b63 | 3 | 5 | 1540.06 | 148.19 | 97.72 | 7.33 |
| **111** | 12993a13 | 3 | 5 | 947.37 | 96.05 | 59.29 | 5.53 |
| **112** | JB Asano | 6 | 5 | 2906.75 | 287.24 | 131.41 | 22.32 |
| **113** | 13031a31 | 4 | 5 | 2131.80 | 237.97 | 99.24 | 11.13 |
| **114** | 13057a41 | 3 | 4 | 776.14 | 86.50 | 55.14 | 2.52 |
| **115** | W140205.1.1 | 4 | 5 | 1759.81 | 201.13 | 105.93 | 4.41 |
| **116** | BR 61192 | 5 | 4 | 1765.42 | 205.94 | 89.05 | 29.42 |
| **117** | 14434p4 | 5 | 7 | 2200.65 | 236.60 | 101.55 | 5.39 |
| **118** | JB Asano | 6 | 6 | 394.03 | 28.98 | 18.28 | 48.77 |
| **119** | JB Asano | 6 | 6 | 860.41 | 62.69 | 44.40 | 35.30 |
| **120** | Floki | 6 | 6 | 2555.09 | 253.80 | 141.03 | 14.57 |
| **121** | Floki | 6 | 6 | 1518.16 | 147.66 | 88.28 | 8.36 |
| **122** | 14639p6z | 5 | 7 | 1751.48 | 172.45 | 80.61 | 2.35 |
| **123** | Floki | 5 | 6 | 527.80 | 54.76 | 24.47 | 8.46 |
| **124** | Floki | 4 | 6 | 3269.32 | 368.26 | 169.72 | 6.31 |
| **125** | Floki | 6 | 7 | 1355.49 | 157.92 | 63.30 | 21.98 |
| **126** | Floki | 5 | 6 | 1161.84 | 123.24 | 53.93 | 10.00 |
| **127** | 13518c5 | 6 | 7 | 333.66 | 29.48 | 19.25 | 10.16 |
| **128** | 13519a4 | 5 | 7 | 461.14 | 66.15 | 20.60 | 21.44 |
| **129** | 13520b6 | 4 | 7 | 216.63 | 14.65 | 12.27 | 5.29 |
| **130** | 13520c2 | 5 | 8 | 3199.04 | 448.61 | 189.84 | 12.47 |
| **131** | Reform | 3 | 6 | 169.34 | 16.98 | 5.76 | 9.67 |
| **132** | Floki | 5 | 7 | 1162.41 | 133.74 | 71.12 | 7.45 |
| **133** | Floki | 6 | 6 | 1292.07 | 142.47 | 76.82 | 5.73 |
| **134** | Floki | 5 | 6 | 890.96 | 107.47 | 48.22 | 20.71 |
| **135** | JB Asano | 3 | 6 | 892.27 | 79.82 | 46.70 | 20.21 |
| **136** | Floki | 6 | 6 | 1853.89 | 285.55 | 94.38 | 10.11 |
| **137** | Floki | 5 | 6 | 494.42 | 52.37 | 23.73 | 8.61 |
| **138** | Floki | 6 | 7 | 337.93 | 32.51 | 20.45 | 7.48 |
| **139** | Floki | 6 | 7 | 2382.38 | 305.22 | 167.53 | 6.47 |
| **140** | Floki | 5 | 7 | 3192.27 | 340.37 | 206.82 | 8.86 |
| **141** | 13804a2 | 7 | 7 | 507.32 | 43.61 | 30.59 | 9.47 |

**Supplementary Table S1.** continued.

| BEO | Variety | FHB 1 | FHB 2 | DON [μg kg$^{-1}$] | DON-3-G [μg kg$^{-1}$] | 3-AcDON [μg kg$^{-1}$] | TeA [μg kg$^{-1}$] |
|---|---|---|---|---|---|---|---|
| **142** | Floki | 8 | 8 | 888.37 | 65.10 | 56.50 | 16.46 |
| **143** | 13810a6 | 7 | 7 | 382.42 | 31.87 | 22.57 | 18.84 |
| **144** | 13992f4 | 4 | 7 | 600.06 | 66.01 | 34.02 | 6.11 |
| **145** | Floki | 5 | 8 | 754.39 | 79.99 | 35.88 | 5.16 |

**Supplementary Table S2.** LC-MS/MS parameters of the analyzed mycotoxins. Detailed are the analyte, its ionization state, precursor mass-to-charge-ratio, two transition product ions mass-to-charge-ratios, respective voltage focusing before Q1, voltage in the collision cell, voltage applied before Q3, and the retention time. Abbreviations in chronological order: AOH = alternariol, AME = alternariol monomethylether, -G = respective glucosides, -S = respective sulfates, ATX I = altertoxin I, TeA = tenuazonic acid, TEN = tentoxin, DON = deoxynivalenol, 3-AcDON = 3-acetyl-DON, NIV = nivalenol, Fus X = fusarenone X, T-2 = T-2 toxin, HT-2 = HT-2 toxin; ZEN = zearalenone, ENN = enniatin, BEA = beauvericin.

| **Analyte** | **Ionization state** | **Precursor ion *m/z*** | **Product ion *m/z*** | **Q1 prebias [V]** | **CE [V]** | **Q3 prebias [V]** | **RT [min]** |
|---|---|---|---|---|---|---|---|
| AOH | $[M – H]^-$ | 257.30 | 213.00/214.85 | 28.0/26.0 | 25.0/23.0 | 34.0/14.0 | 9.45 |
| $[^{2}H_4]$-AOH | $[M – H]^-$ | 261.30 | 217.00/218.85 | 28.0/26.0 | 25.0/23.0 | 34.0/14.0 | 9.45 |
| AOH-3G | $[M – H]^-$ | 419.10 | 256.15/255.10 | 30.0/16.0 | 33.0/44.0 | 26.0/26.0 | 7.22 |
| AOH-9G | $[M – H]^-$ | 419.10 | 256.15/255.10 | 30.0/16.0 | 33.0/44.0 | 26.0/26.0 | 8.82 |
| AOH-3S | $[M – H]^-$ | 337.10 | 257.15/213.10 | 24.0/24.0 | 22.0/40.0 | 24.0/18.0 | 7.49 |
| AOH-9S | $[M – H]^-$ | 337.10 | 257.15/213.10 | 24.0/24.0 | 22.0/40.0 | 24.0/18.0 | 8.77 |
| AME | $[M – H]^-$ | 271.25 | 256.00/255.10 | 14.0/34.0 | 23.0/28.0 | 10.0/26.0 | 12.76 |
| [2H4]-AME | $[M – H]^-$ | 275.25 | 260.00/259.10 | 14.0/34.0 | 23.0/28.0 | 10.0/26.0 | 12.76 |
| AME-3G | $[M – H]^-$ | 433.30 | 270.20/271.20 | 16.0/12.0 | 33.0/26.0 | 18.0/20.0 | 11.20 |
| AME-3S | $[M – H]^-$ | 351.20 | 271.20/256.15 | 12.0/12.0 | 23.0/35.0 | 26.0/24.0 | 10.84 |
| ATX I | $[M – H]^-$ | 351.20 | 315.00/333.05 | 26.0/26.0 | 17.0/12.0 | 18.0/36.0 | 11.20 |
| TeA | $[M – H]^-$ | 196.40 | 111.95/139.00 | 22.0/22.0 | 25.0/19.0 | 34.0/26.0 | 4.65 |
| $[^{13}C_6, ^{15}N]$-TeA | $[M – H]^-$ | 203.40 | 112.95/142.00 | 22.0/22.0 | 25.0/19.0 | 34.0/26.0 | 4.65 |
| TEN | $[M – H]^-$ | 413.40 | 141.05/271.30 | 14.0/14.0 | 23.0/20.0 | 12.0/16.0 | 11.78 |
| DON | $[M + H]^+$ | 297.15 | 249.20/231.15 | -14.0/-18.0 | -11.0/-12.0 | -18.0/-26.0 | 5.75 |
| $[^{15}C_{13}]$-DON | $[M + H]^+$ | 312.15 | 263.20/245.15 | -14.0/-18.0 | -11.0/-12.0 | -18.0/-26.0 | 5.75 |
| DON-3G | $[M – H]^-$ | 457.30 | 427.05/255.35 | 22.0/16.0 | 17.0/27.0 | 44.0/30.0 | 6.05 |
| $[^{13}C_{21}]$-DON-3G | $[M – H]^-$ | 478.15 | 447.05/261.30 | 22.0/16.0 | 17.0/27.0 | 44.0/30.0 | 6.05 |
| 3-AcDON | $[M – H]^-$ | 339.10 | 231.25/175.20 | -16.0/-16.0 | -13.0/-25.0 | -26.0/-20.0 | 8.73 |
| $[^{13}C_{17}]$-3-AcDON | $[M – H]^-$ | 356.10 | 245.25/186.20 | -16.0/-16.0 | -13.0/-25.0 | -26.0/-20.0 | 8.73 |
| NIV | $[M – H]^-$ | 311.20 | 281.15/191.20 | 22.0/12.0 | 11.0/21.0 | 16.022.0 | 4.65 |
| Fus X | $[M + H]^+$ | 372.15 | 355.25/337.20 | -18.0/-18.0 | -9.0/-13.0 | -18.0/-24.0 | 7.40 |
| T-2 | $[M + NH_4]^+$ | 484.35 | 305.20/215.30 | -26.0/-28.0 | -15.0/-21.0 | -22.0/-24.0 | 12.13 |
| $[^{13}C_4]$-T2 | $[M + NH_4]^+$ | 488.35 | 307.15/216.25 | -26.0/-28.0 | -15.0/-21.0 | -22.0/-24.0 | 12.13 |
| HT-2 | $[M + NH_4]^+$ | 442.20 | 263.25/215.15 | -22.0/-22.0 | -14.0/-14.0 | -10.0/-24.0 | 11.57 |
| $[^{13}C_{22}]$-HT2 | $[M + NH_4]^+$ | 464.20 | 278.25/200.15 | -22.0/-22.0 | -14.0/-14.0 | -10.0/-24.0 | 11.57 |
| ZEN | [M – H]- | 413.40 | 141.05/271.30 | 14.0/14.0 | 23.0/20.0 | 12.0/16.0 | 12.37 |
| ENN A1 | $[M + NH_4]^+$ | 685.45 | 210.25/100.30 | -20.0/-16.0 | -29.0/-55.0 | -14.0/-22.0 | 14.70 |
| $[^{15}N_3]$-ENN A1 | $[M + NH_4]^+$ | 688.45 | 211.25/101.30 | -20.0/-16.0 | -29.0/-55.0 | -14.0/-22.0 | 14.70 |
| ENN A | $[M + NH_4]^+$ | 699.45 | 210.20/100.30 | -16.0/-16.0 | -33.0/-55.0 | -22.0/-24.0 | 14.85 |
| ENN B1 | $[M + NH_4]^+$ | 671.35 | 196.20/210.25 | -16.0/-16.0 | -36.0/-32.0 | -14.0/-14.0 | 14.65 |
| ENN B | $[M + NH_4]^+$ | 657.45 | 196.20/86.20 | -18.0/-26.0 | -32.0/-64.0 | -14.0/-18.0 | 14.30 |
| BEA | $[M + NH_4]^+$ | 801.50 | 244.20/134.20 | -18.0/-18.0 | -33.0/-55.0 | -18.0/-14.0 | 14.50 |

**Supplementary Table S3.** Games–Howell post-hoc comparisons of DON concentrations between FHB severity classes following Welch's ANOVA. Shown are mean differences (class B – class A), 95% confidence intervals, adjusted p-values, and significance levels (ns = p ≥ 0.05; * = p < 0.05; ** = p < 0.01; *** = p < 0.001; **** = p < 0.0001).

| Variable | FHB class (A) | FHB class (B) | Mean difference (µg/kg) | 95% CI lower | 95% CI upper | Adjusted p-value | Significance |
|---|---|---|---|---|---|---|---|
| DON | 1 | 2 | 510.35631 | 107.60586 | 913.1068 | 5.00E-03 | ** |
| DON | 1 | 3 | 672.32825 | 281.37513 | 1063.2814 | 3.20E-05 | **** |
| DON | 1 | 4 | 1404.3224 | 885.13164 | 1923.5131 | 1.91E-10 | **** |
| DON | 1 | 5 | 1357.43 | 937.46232 | 1777.3976 | 8.48E-12 | **** |
| DON | 1 | 6 | 1431.8312 | 785.37601 | 2078.2863 | 5.89E-07 | **** |
| DON | 1 | 7 | 896.07621 | 47.666 | 1744.4864 | 3.50E-02 | * |
| DON | 1 | 8 | 1277.049 | -2400.536 | 4954.6339 | 5.22E-01 | ns |
| DON | 2 | 3 | 161.97194 | -367.08744 | 691.0313 | 9.80E-01 | ns |
| DON | 2 | 4 | 893.96605 | 266.85326 | 1521.0789 | 6.74E-04 | *** |
| DON | 2 | 5 | 847.07366 | 296.73864 | 1397.4087 | 1.74E-04 | *** |
| DON | 2 | 6 | 921.47486 | 190.6336 | 1652.3161 | 5.00E-03 | ** |
| DON | 2 | 7 | 385.7199 | -512.21685 | 1283.6567 | 8.31E-01 | ns |
| DON | 2 | 8 | 766.69265 | -2782.2581 | 4315.6434 | 8.68E-01 | ns |
| DON | 3 | 4 | 731.99412 | 110.51535 | 1353.4729 | 1.00E-02 | ** |
| DON | 3 | 5 | 685.10172 | 141.63069 | 1228.5727 | 4.00E-03 | ** |
| DON | 3 | 6 | 759.50292 | 33.11387 | 1485.892 | 3.40E-02 | * |
| DON | 3 | 7 | 223.74796 | -671.25362 | 1118.7495 | 9.89E-01 | ns |
| DON | 3 | 8 | 604.72071 | -2949.5408 | 4158.9822 | 9.46E-01 | ns |
| DON | 4 | 5 | -46.89239 | -686.39241 | 592.6076 | 1.00E+00 | ns |
| DON | 4 | 6 | 27.5088 | -769.2446 | 824.2622 | 1.00E+00 | ns |
| DON | 4 | 7 | -508.24615 | -1452.0643 | 435.572 | 6.50E-01 | ns |
| DON | 4 | 8 | -127.27341 | -3587.3009 | 3332.7541 | 1.00E+00 | ns |
| DON | 5 | 6 | 74.4012 | -666.78404 | 815.5864 | 1.00E+00 | ns |
| DON | 5 | 7 | -461.35376 | -1366.138 | 443.4305 | 6.89E-01 | ns |
| DON | 5 | 8 | -80.38101 | -3613.6292 | 3452.8672 | 1.00E+00 | ns |
| DON | 6 | 7 | -535.75495 | -1539.2199 | 467.71 | 6.72E-01 | ns |
| DON | 6 | 8 | -154.78221 | -3529.1954 | 3219.631 | 1.00E+00 | ns |
| DON | 7 | 8 | 380.97274 | -2904.1023 | 3666.0478 | 9.97E-01 | ns |
| **Welch's ANOVA** | | **Variable** | **Sample size** | **F-value** | **df (between groups)** | **df (within groups)** | **p-value** |
| | | DON | 290 | 30.81 | 7 | 41.78 | $1.53 \times 10^{-14}$ |

**Supplementary Table S4.** UPLC-parameters and gradient for metabolomics measurements.

| Parameter | Settings | |
|---|---|---|
| Column | ACQUITY UPLC BEH C18 Column 130 Å, 1.7 µm, 2.1 mm x 100 mm | |
| Mobile phase A | Ultrapure $H_2O$ (Milli-Q) + 0.1%Formic acid (Honeywell Chromasolv LC-MS grade) | |
| Mobile phase B | Acetonitrile (Honeywell Chromasolv LC-MS grade) + 0.1%Formic acid | |
| Flow rate | 0.4 mL/min | |
| Gradient | Time [min] | B.Conc [%] |
| | 0.00 | 5 |
| | 1.20 | 5 |
| | 3.70 | 50 |
| | 11.50 | 95.5 |
| | 15.25 | 95.5 |
| | 15.75 | 5.0 |
| | 19.00 | 5.0 |
| Oven temperature | 40 °C | |
| Injection volume | 5 µL | |

**Supplementary Table S5.** ToF-MS and SWATH parameters for metabolomics measurements.

| MS-parameters | | |
|---|---|---|
| Method duration [min] | 15.25 | |
| Total scan time [sec] | 0.82 | |
| Estimated cycles | 1117 | |
| Source | Turbulon Spray | |
| Curtain gas [psi] | 30 | |
| Ion source gas1 [psi] | 45 | |
| Ion source gas 2 [psi] | 45 | |
| Temperature [°C] | 500 | |
| Scan type | SWATH | |
| CAD gas | 7 | |
| ToF-MS1 | Positive | Negative |
| Spray voltage [V] | 5500 | -4500 |
| Time bins to sum | 4 | 4 |
| ToF-MS1 start mass [Da] | 100 | 100 |
| ToF-MS1 stop mass [Da] | 1500 | 1500 |
| Accumulation time [sec] | 0.1 | 0.1 |
| Declustering potential | 50 | -80 |
| DP spread [V] | 0 | 0 |
| Collision energy [V] | 5 | -5 |
| Collision energy spread [V] | 0 | |
| QJet amplitude [V] | 193.788 | 193.788 |
| ToF-MS2 | SWATH | |
| ToF-MS2 start mass [Da] | 120 | 120 |
| ToF-MS2 stop mass [Da] | 1500 | 1500 |
| Accumulation time [sec] | 0.03 | 0.03 |
| Dynamic collision energy | False | False |
| QJet amplitude [V] | 200.728 | 200.728 |
| Charge state | 1 | 1 |

| SWATH-parameters | |
|---|---|
| Collision energy spread [V] | 15 |
| Time bins to sum | 8 |
| Declustering potential [V] | 80 (positive) and -50 (negative) |
| Declustering potential spread [V] | 0 |

| Positive | | | Negative | | |
|---|---|---|---|---|---|
| Start mass [Da] | Stop mass [Da] | CE [V] | Start mass [Da] | Stop mass [Da] | CE [V] |
| 99.5 | 210.7 | 12 | 99.5 | 237.7 | 12 |
| 209.7 | 245.8 | 12 | 236.7 | 271.8 | 12 |
| 245.8 | 271.5 | 15 | 270.8 | 299.1 | 15 |
| 270.5 | 293.3 | 15 | 298.1 | 324.5 | 15 |
| 292.3 | 312.3 | 15 | 323.5 | 349.8 | 15 |
| 311.3 | 330.4 | 20 | 348.8 | 374.2 | 20 |
| 329.4 | 349.4 | 20 | 373.2 | 408.3 | 20 |
| 348.4 | 371.2 | 20 | 407.3 | 446.4 | 20 |

**Supplementary Table S5.** continued.

| | | | | | |
|---|---|---|---|---|---|
| 370.2 | 398.8 | 25 | 445.4 | 478.5 | 25 |
| 397.8 | 435.8 | 25 | 477.5 | 504.9 | 25 |
| 434.8 | 471.0 | 25 | 503.9 | 529.2 | 25 |
| 470.0 | 503.3 | 30 | 528.2 | 556.5 | 30 |
| 502.3 | 536.5 | 30 | 555.5 | 584.8 | 30 |
| 535.5 | 569.8 | 30 | 583.8 | 614.1 | 30 |
| 568.8 | 607.8 | 35 | 613.1 | 651.1 | 35 |
| 606.8 | 655.3 | 35 | 650.1 | 696.0 | 35 |
| 654.3 | 704.7 | 35 | 695.0 | 757.4 | 35 |
| 703.7 | 777.9 | 40 | 756.4 | 849.0 | 40 |
| 776.9 | 947.9 | 40 | 848.0 | 1025.5 | 40 |
| 946.9 | 1498.9 | 40 | 1024.5 | 1499.5 | 40 |

**Supplementary Table S6.** MS-DIAL-parameters for metabolomics data processing.

| Parameter | settings | |
|---|---|---|
| | Data collection | |
| MS1 tolerance | 0.005 Da | |
| MS2 tolerance | 0.01 Da | |
| Retention time range | 3.0 - 12.0 min | |
| MS1 range | 100 - 1500 Da | |
| MS2 range | 20 - 1500 Da | |
| Maximum charges | 2 | |
| | Peak detection | |
| Minimum peak hight | 2000 amplitude | |
| Mass slice width | 0.05 Da | |
| Smoothing method | Savitzky-Golay filter | |
| Smoothing level | 5 scans | |
| Minimum peak width | 5 scans | |
| Exclusion list (+- 0.005 Da) | Positive: *m/z* 132.9049, 266.1598, 354.2122, 442.2647, 609.2807, 829.5393, 961.4370, 1446,7322 | Negative: *m/z* 68.9996, 112.9856, 154.9738, 204.9706, 248,96040, 384.9352, 520.9100, 656.8848, 792.8596, 928.8344, 1064.8092, 1200.7841, 1336.7589, 1472.7337 |
| | Spectrum deconvolution | |
| Sigma window value | 0.8 | |
| MS/MS abundance cutoff | 120 amplitude | |
| Explude after precurser ion | yes | |
| Keep isotopic ions | yes | |
| Adduct ions | | |
| | Positive: $[M+H]^+$, $[M+Na]^+$, $[M+K]^+$, $[M+ACN+H]^+$, $[M+H-H_2O]^+$, $[2M+H]^+$, $[2M+Na]^+$, $[M+2H]^{2+}$ | Negative: $[M-H]^-$, $[M-H_2O-H]^-$, $[M-FA]^-$, [2M-H]-, $[2M+FA-H]^-$, $[M-2H]^{2-}$ |
| | Alignment parameter | |
| Reference files | QC measurements | |
| Retention time tolerance | 0.1 min | |
| MS1 tolerance | 0.005 Da | |
| Retention time factor | 0.6 | |
| MS1 factor | 0.4 | |
| Peak count filter | 1% | |
| Sample max / blank avg filter | 10-fold change | |
| | .mgf file export | |
| MGF export | Merge over all samples (MS1 0.005 Da) | |
| | MS2 0.01 Da or 20 ppm | |
| | Cosine threshold 0.6 | |
| | Signal count threshold 30% | |
| | Data transformation | |
| Zero-filling | Random number between $(avg_{\text{minimum intensity in each sample}} - \sigma \pm \sigma)$ | |
| Normalization | z-score | |

**Supplementary Table S7.** SIRIUS Tandem-mass spectra characterization parameters.

| Parameter | settings |
|---|---|
| | SIRIUS |
| Molecular formula identification | $C_{\infty}H_{\infty}$ $N_8$ $O_{\infty}$ $S_3$ $P_1$ |
| Ionizations and Integer Linear Programming (ILP) | Default $[M+H]^+$, $[M+Na]^+$, $[M+K]^+$, $[M+ACN+H]^+$, $[M+H-H_2O]^+$, $[2M+H]^+$, $[M+2H]^{2+}$ |
| SCI:FingerID fingerprint prediction | $MS^2$ Mass accuracy 10 ppm |
| CSI:FingerID structure database search | Bio Database, KEGG, Biocyc, KNApSAcK, CHEBI, Maconda, COCONUT, NORMAN, EcoCycMine, Natural Products, GNPS, Plantcyc, HMDB, YMDB |
| CANOPUS compound class prediction | Main class: Level 5<br>Natural Product Class: class |

**Supplementary Table S8.** Statistical values describing the PCA and OPLS models.

| Component | Values | | | | | |
|---|---|---|---|---|---|---|
| | **Principal Component Analysis (PCA)** | | | | | |
| | R2X | R2X(cum) | | | | |
| Principal Comp. 1 | 0.127 | 0.127 | | | | |
| Principal Comp. 2 | 0.061 | 0.188 | | | | |
| Principal Comp. 3 | 0.037 | 0.225 | | | | |
| Principal Comp. 4 | 0.030 | 0.255 | | | | |
| Principal Comp. 5 | 0.027 | 0.282 | | | | |
| | **Orthogonal Projection to Latent Structures (OPLS)** | | | | | |
| | R2X | R2X(cum) | R2Y | R2Y(cum) | Q2 | Q2(cum) |
| Predictive Comp. 1 | 0.023 | 0.023 | 0.752 | 0.752 | 0.699 | 0.699 |
| Orthogonal Comp. 2 | 0.1 | 0.123 | 0.111 | 0.111 | 0.124 | 0.124 |
| Orthogonal Comp. 3 | 0.079 | 0.202 | 0.053 | 0.164 | 0.057 | 0.181 |
| Orthogonal Comp. 4 | 0.026 | 0.228 | 0.051 | 0.215 | 0.054 | 0.235 |
| Orthogonal Comp. 5 | 0.022 | 0.250 | 0.014 | 0.228 | 0.012 | 0.247 |
| | | 0.250 | | **0.98** | | **0.946** |

**Supplementary Table S9.** Tandem-mass spectrometric data of the most significant features _positively_ correlated with the DON concentration quantified in the wheat grain samples. The fragments clearly indicating a sesquiterpene core structure are highlighted in bold. Duplicates due to ion-source fragmentation or adduct formation are faded in grey.

| VIP | ID | *m/z* | Formula | Fragmentation spectrum *m/z* (formula, rel. intensity) |
|---|---|---|---|---|
| | | | Positively correlating with the DON-concentration | |
| 6.26 | 1506 | 265.1801 | $C_{16}H_{26}O_4$ [-$H_2O$+H]$^+$ | 219.174(C15H22O, 0.11), 201.1646(C15H20, 0.24), 175.1482(C13H18, 0.63), 173.1318(C13H16, 0.1), 159.1172(C12H14, 0.17), 147.1172(C11H14, 0.13), 145.1017(C11H12, 0.43), **135.1174**(C10H14, 0.13), 123.0806(C8H10O, 0.03), **121.1015**(C9H12, 0.28), **119.0856**(C9H10, 1), **109.1015**(C8H12, 0.27), **107.0863**(C8H10, 0.38), 105.0697(C8H8, 0.62), **95.0858**(C7H10, 0.37), 93.0704(C7H8, 0.33), 91.0545(C7H6, 0.25), **81.0696**(C6H8, 0.07), 67.0543(C5H6, 0.04), 55.0543(C4H6, 0.06), 43.0176(C2H2O, 0.04) |
| 6.23 | 603 | 201.1641 | $C_{15}H_{22}O$ [-$H_2O$+H]$^+$ | 159.1179(C12H14, 0.09), 145.1019(C11H12, 0.2), 143.0949(C11H10, 0.13), 142.0782(C11H9, 0.14), 129.0703(C10H8, 0.28), **119.0857**(C9H10, 0.24), **115.0548(**C9H6, 0.32), **109.1023**(C8H12, 0.08), **105.0704**(C8H8, 0.63), 103.0549(C8H6, 0.21), **95.0858**(C7H10, 0.13), 91.0549(C7H6, 1), 69.0701(C5H8, 0.01), 66.0444(C5H5, 0.15), 41.039(C3H4, 0.49) |
| 6.21 | 811 | 219.1742 | $C_{15}H_{22}O$ [+H]$^+$ | 219.1738(C15H22O, 0.13), 201.1642(C15H20, 0.17), 177.1639(C13H20, 0.11), 175.1485(C13H18, 0.2), 159.1175(C12H14, 0.35), 145.1014(C11H12, 0.7), 143.0863(C11H10, 0.11), **135.1173**(C10H14, 0.07), **133.1014**(C10H12, 0.3), 131.0854(C10H10, 0.24), 129.0704(C10H8, 0.17), **121.102**(C9H12, 0.18), **119.0858**(C9H10, 1), **109.1012**(C8H12, 0.28), **107.086**(C8H10, 0.26), **105.0702**(C8H8, 0.89), 103.0557(C8H6, 0.11), **95.086**(C7H10, 0.41), 91.0546(C7H6, 0.38), **81.0703**(C6H8, 0.13), **69.0702**(C5H8, 0.1), 67.0552(C5H6, 0.15) |
| 6.19 | 1265 | 251.1647 | $C_{15}H_{22}O_3$ [+H]$^+$ | 187.1466(C14H18, 0.3), 175.1118(C12H14O, 0.3), 159.1178(C12H14, 0.55), 145.1013(C11H12, 0.5), 143.0866(C11H10, 0.3), 131.0856(C10H10, 0.61), **121.1017**(C9H12, 0.53), **119.0856**(C9H10, 0.63), **109.1013**(C8H12, 0.58), 105.07(C8H8, 1), **95.086**(C7H10, 0.76), **93.0705**(C7H8, 0.81), **81.0705**(C6H8, 0.4) |
| 6.09 | 2313 | 300.2172 | $C_{16}H_{29}NO_4$ [+H]$^+$ | 265.1802(C16H24O3, 0.11), 219.1744(C15H22O, 0.51), 177.1645(C13H20, 0.33), 175.1484(C13H18, 1), 151.1112(C10H14O, 0.51), **149.13227**(C11H16, 0.06), 147.1166(C11H14, 0.1), 145.1022(C11H12, 0.2), 137.0961(C9H12O, 0.05), **133.1016**(C10H12, 0.18), 125.0962(C8H12O, 0.02), 123.1168(C9H14, 0.07), **121.1014**(C9H12, 0.29), **119.0861**(C9H10, 0.44), 111.0808(C7H10O, 0.03), **109.101**(C8H12, 0.06), 99.0811(C6H10O, 0.01), 97.1015(C7H12, 0.11), **95.0857**(C7H10, 0.35), **93.0703**(C7H8, 0.07), 79.0546(C6H6, 0.05), 71.0857(C5H10, 0.01), 55.0544(C4H6, 0.05) |
| 6.05 | 1030 | 235.1692 | $C_{15}H_{22}O_2$ [+H]$^+$ | 175.1494(C13H18, 0.2), 161.1327(C12H16, 0.23), 159.1175(C12H14, 0.33), 147.1175(C11H14, 0.23), 145.1019(C11H12, 0.53), 143.0869(C11H10, 0.23), **135.117**(C10H14, 0.2), **133.1015**(C10H12, 0.32), 131.0861(C10H10, 0.3), 129.0704(C10H8, 0.42), **121.1018**(C9H12, 0.35), **119.0857**(C9H10, 0.64), 117.07(C9H8, 0.28), 111.0446(C6H6O2, 0.36), **109.1008**(C8H12, 0.37), **107.0859**(C8H10, 0.51), **105.0707**(C8H8, 1), 101.0601(C5H8O2, 0.33), 95.086(C7H10, 0.54), **93.0703**(C7H8, 0.38), 91.055(C7H6, 0.77), 83.0858(C6H10, 0.34), **81.0706(**C6H8, 0.33), 79.0546(C6H6, 0.32), 67.0547(C5H6, 0.43), 59.0711(C5H8, 0.28), 55.0547(C4H6, 0.62), 43.0184(C2H2O, 0.54) |
| 6.04 | 854 | 221.1900 | $C_{15}H_{24}O$ [+H]$^+$ | 221.1912(C15H24O, 0.09), 203.1809(C15H22, 0.22), 177.1646(C13H20, 0.34), 175.149(C13H18, 0.07), 161.1333(C12H16, 0.18), **149.1343**(C11H16, 0.06), 147.118(C11H14, 0.37), 145.1022(C11H12, 0.05), **135.1179**(C10H14, 0.15), **133.1022**(C10H12, 0.35), 131.0869(C10H10, 0.02), 123.1182(C9H14, 0.13), **121.102**(C9H12, 0.55), **119.0863**(C9H10, 0.72), 117.0706(C9H8, 0.07), **115.0552**(C9H6, 0.02), **109.102**(C8H12, 0.5), **107.0861**(C8H10, 0.78), **105.0707**(C8H8, 1), 103.0549(C8H6, 0.07), **95.0864**(C7H10, 0.71), **93.071**(C7H8, 0.4), 91.0549(C7H6, 0.54), 83.066(C6H10, 0.05), **81.071**(C6H8, 0.15), 79.0551(C6H6, 0.05), **69.0711**(C5H8, 0.16), 67.055(C5H6, 0.1), 57.0705(C4H8, 0.04), 55.055(C4H6, 0.13), 43.0547(C3H6, 0.11) |

**Supplementary Table S9.** continued.

| VIP | ID | *m/z* | Formula | Fragmentation spectrum *m/z*(formula, rel. intensity) |
|---|---|---|---|---|
| 6.04 | 1008 | 233.1535 | $C_{15}H_{20}O_2$ [+H]$^+$ | 175.148(C13H18, 0.37), 159.1172(C12H14, 0.44), 157.1018(C12H12, 0.31), 155.0868(C12H10, 0.29), 147.1163(C11H14, 0.3), 145.1009(C11H12, 0.5), 143.0855(C11H10, 0.44), **133.1023**(C10H12, 0.46), 131.0859(C10H10, 0.57), 123.0809(C8H10O, 0.29), **119.0863**(C9H10, 1), **109.1021**(C8H12, 0.54), 107.0861(C8H10, 0.47), **105.07**(C8H8, 0.8505), **95.086**(C7H10, 0.86), **93.07**(C7H8, 0.567), 91.0547(C7H6, 0.78), **81.07**(C6H8, 0.34), 67.0554(C5H6, 0.29), 55.0549(C4H6, 0.36) |
| 6.03 | 620 | 203.1796 | $C_{15}H_{24}O$ [-$H_2O$+H]$^+$ | 203.1789(C15H22, 0.13), 161.1339(C12H16, 0.14), 147.1171(C11H14, 0.2), 143.0844(C11H10, 0.18), **133.1013**(C10H12, 0.42), 131.0856(C10H10, 0.16), 129.0696(C10H8, 0.24), **121.1014**(C9H12, 0.27), **119.0859**(C9H10, 0.74), 117.0702(C9H8, 0.27), **115.0542**(C9H6, 0.34), **109.1012**(C8H12, 0.32), **107.0854**(C8H10, 0.36), **105.0699**(C8H8, 0.8), 103.0545(C8H6, 0.26), **95.086**(C7H10, 0.42), 91.0545(C7H6, 0.98), **81.0702**(C6H8, 0.17), 77.0386(C6H4, 0.21), **69.0704**(C5H8, 0.16), 67.0539(C5H6, 0.23), 57.071(C4H8, 0.12), 55.0545(C4H6, 1), 41.0386(C3H4, 0.72) |
| 6.02 | 1062 | 237.1851 | $C_{15}H_{24}O_2$ [+H]$^+$ | 237.1852(C15H24O2, 0.05), 219.1749(C15H22O, 0.14), 159.1175(C12H14, 0.02), 149.097(C10H12O, 0.09), 145.1012(C11H12, 0.02), **135.0804**(C9H10O, 1), **133.1019**(C10H12O, 0.01), **121.0651**(C8H8O, 0.4), **119.0868**(C9H10, 0.05), **115.0551**(C9H6, 0.04), **109.1021**(C8H12, 0.05), **107.0862**(C8H10, 0.06), **105.0704**(C8H8O, 0.18), **95.0866**(C7H10, 0.03), 91.055(C7H6, 0.08), **81.0704**(C6H8, 0.01), **69.0701**(C5H8, 0.05), 55.0548(C4H6, 0.04) |
| 6.00 | 1010 | 233.1543 | $C_{15}H_{20}O_2$ [+H]$^+$ | 173.1334(C13H16, 0.14), **133.1013**(C10H12, 0.17), 131.0858(C10H10, 0.19), **121.1019**(C9H12, 0.2), **119.0492**(C8H6O, 0.75), 117.0701(C9H8, 0.21), **115.0536**(C9H6, 0.14), 111.0438(C6H6O2, 0.22), **109.1009**(C8H12, 0.6), **105.0705**(C8H8, 0.66), 97.065(C6H8O, 0.27), **95.0855**(C7H10, 0.45), 91.0546(C7H6, 1), 83.0499(C5H6O, 0.14), **81.0699**(C6H8O, 0.46), 79.0546(C6H6O2, 0.16), 67.0547(C5H6, 0.23), 57.0704(C4H8, 0.22), 55.0543(C4H6, 0.48), 43.0181(C2H2O, 0.64) |
| 5.94 | 339 | 177.1639 | $C_{13}H_{20}$ [+H]$^+$ | 161.1339(C12H16, 0.14), 147.1171(C11H14, 0.2), 143.0844(C11H10, 0.18), **133.1013**(C10H12, 0.42), 131.0856(C10H10, 0.16), 129.0696(C10H8, 0.24), **121.1014**(C9H12, 0.27), **119.0859**(C9H10, 0.74), 117.0702(C9H8, 0.27), **115.0542**(C9H6, 0.34), 109.1012(C8H12, 0.32), **107.0854**(C8H10, 0.36), 105.0699(C8H8, 0.8), 103.0545(C8H6, 0.26), **95.086**(C7H10, 0.43), 91.0545(C7H6, 0.98), **81.0702**(C6H8, 0.17), 77.0386(C6H4, 0.21), 69.0704(C5H8, 0.16), 67.0539(C5H6, 0.23), 57.071(C4H8, 0.12), 55.0545(C4H6, 1), 41.0386(C3H4, 0.72) |
| 5.89 | 1911 | 283.1901 | $C_{16}H_{26}O_4$ [+H]$^+$ | 219.1749(C15H22O, 0.46), 201.1637(C15H20, 0.45), 175.148(C13H18, 0.07), 163.1126(C11H14O, 0.06), 161.1325(C12H16, 0.4), 159.1172(C12H14, 0.29), 145.1013(C11H12, 0.01), 131.0859(C10H10, 0.51), **121.1018**(C9H12, 0.06), 119.0854(C9H10, 0.25), 117.07(C9H8, 0.22), 111.0805(C7H10O, 0.01), **109.1012**(C8H12, 0.44), **105.0699**(C8H8, 1), **95.0858**(C7H10O, 0.3), **93.0702**(C7H8, 0.67), 43.0181(C2H2O, 0.13) |
| 5.79 | 2387 | 305.1731 | $C_{16}H_{26}O_4$ [+Na]$^+$ | 219.1748(C15H22O, 0.37), 201.1649(C15H20, 0.25), 177.1635(C13H20, 0.27), 175.1489(C13H18, 0.54), 165.1258(C11H16O, 0.16), 163.1131(C11H14O, 0.17), 161.1336(C12H16, 0.16), 159.1176(C12H14, 0.17), 151.1122(C10H14O, 0.35), 137.0965(C9H12O, 0.29), **133.1015**(C10H12, 0.31), **121.1018**(C9H12, 0.32), **119.0862**(C9H10, 0.43), **109.1013**(C8H12, 0.55), **107.0861**(C8H10, 0.39), **105.0699**(C8H8, 0.34), 99.0807(C6H10O, 0.19), 97.1011(C7H12, 0.25), **95.0863**(C7H10, 0.86), 95.0495(C6H6O, 1), **81.0708**(C6H8, 0.32), 71.0854(C5H10, 0.23), 67.055(C5H6, 0.32), 55.0544(C4H6, 0.35), 43.0545(C3H6, 0.33) |
| 5.74 | 2317 | 300.2532 | $C_{18}H_{32}O_3$ [-$H_2O$+H]$^+$ | - |
| 5.73 | 809 | 219.1741 | $C_{15}H_{24}O_2$ [-$H_2O$+H]$^+$ | 219.1746(C15H22O, 0.12), **149.0962**(C10H12O, 0.07), **135.0807**(C9H10O, 1), 123.0805(C8H10O, 0.02), **121.065**(C8H8O, 0.35), **119.0862**(C9H10, 0.02), **109.1014**(C8H12, 0.07), **107.0859**(C8H10, 0.05), **105.0699**(C8H8O, 0.1), **95.0861**(C7H10, 0.04), 91.0545(C7H6, 0.09), **69.0709**(C5H8, 0.07), 55.055(C4H6, 0.03) |

**Supplementary Table S9.** continued.

| VIP | ID | *m/z* | Formula | Fragmentation spectrum *m/z*(formula, rel. intensity) |
|---|---|---|---|---|
| 5.7 | 499 | 193.1587 | $C_{13}H_{20}O$ [+H]$^+$ | **119.0863**(C9H10, 0.23), **115.0555**(C9H6, 0.2), **107.086**(C8H10, 0.18), **105.0701**(C8H8, 0.21), **95.0862**(C7H10, 0.21), 91.0546(C7H6, 0.81), 83.0491(C5H6O, 0.16), **69.0704**(C5H7, 0.17), 65.0395(C5H4, 0.25), 59.0491(C3H6O, 0.28), 57.0702(C4H8, 0.16), 55.0544(C4H6, 1.00), 41.0389(C3H4, 0.45) |
| 5.69 | 2319 | 300.2535 | $C_{17}H_{33}NO_3$ [+H]$^+$ | 46.0657(C2H7N, 1.00) |
| 5.69 | 324 | 175.1485 | $C_{13}H_{18}$ [+H]$^+$ | 145.1024(C11H12, 0.09), 143.0853(C11H10, 0.16), 128.0622(C10H7, 0.27), **121.1013**(C9H12, 0.07), **115.0548**(C9H6, 0.19), **107.0862**(C8H10, 0.12), **105.0704**(C8H8, 0.49), 103.0549(C8H6, 0.15), **95.0861**(C7H10, 0.14), 91.0546(C7H6, 1), **81.0701**(C6H8, 0.09), 79.0547(C6H6, 0.19), **69.0705**(C5H8, 0.09), 67.0551(C5H6, 0.18), 55.0547(C4H6, 0.58), 57.0701(C4H8, 0.15), 41.0392(C3H4, 0.29) |
| 5.64 | 501 | 193.1588 | $C_{13}H_{20}O$ [+H]$^+$ | **135.1181**(C10H14, 0.17), 117.0703(C9H8, 0.11), **115.0546**(C9H6, 0.11), **109.102**(C8H12, 0.09), **107.0863**(C8H10, 0.16), **105.0703**(C8H8, 0.24), **95.0858**(C7H10, 0.1611), 91.0545(C7H6, 1), **81.0708**(C6H8, 0.09), 79.0547(C6H6, 0.11), **69.0695**(C5H8, 0.09), 55.0546(C4H6, 0.41), 41.0385(C3H4, 0.34) |

**Supplementary Table S10.** Tandem-mass spectrometric data of the most significant features _negatively_ correlated with the DON concentration quantified in the wheat grain samples. The fragments indicating a hexosyl-neutral loss (-$C_6H_{10}O_5$, 162.0528) are highlighted in bold.

| VIP | ID | *m/z* | Formula | Fragmentation spectrum *m/z*(formula, rel. intensity) |
|---|---|---|---|---|
| | | | Negatively correlating with the DON-concentration | |
| 3.47 | 6775 | 501.3061 | $C_{26}H_{44}O_9$ [+H]$^+$ | 483.2933(**C26H42O8**, 0.4), 321.2418(**C20H32O3**, 0.43), 305.2476(C20H32O2, 0.44), 303.2312(C20H30O2, 0.37), 165.0912(C10H12O2, 0.41), 139.1121(C9H14O, 0.36), 137.1327(C10H16, 0.37), 135.1164(C10H14, 0.35), 123.1178(C9H14, 0.49), 109.065(C7H8O, 1), 95.0496(C6H6O, 0.64), 93.07(C7H8, 0.52) |
| 3.38 | 6774 | 501.3060 | $C_{26}H_{44}O_9$ [+H]$^+$ | 483.2933(**C26H42O8**, 0.4), 321.2431(**C20H32O3**, 0.27), 307.2635(C20H34O2, 0.55), 305.2475(C20H32O2, 0.22), 303.2316(C20H30O2, 0.66), 293.2121(C18H28O3, 0.23), 277.217(C18H28O2, 0.56), 207.174(C14H22O, 0.22), 189.1646(C14H20, 0.23), 183.102(C10H14O3, 0.25), 179.106(C11H14O2, 0.2124), 177.0919(C11H12O2, 0.22), 175.1484(C13H18, 0.22), 165.0914(C10H12O2, 0.59), 161.0972(C11H12O2, 0.25), 151.1121(C10H14O, 0.26), 149.1326(C11H16, 0.35), 147.1177(C11H14O2, 0.24), 135.1182(C10H14, 0.6), 133.1021(C10H12O2, 0.38), 127.0397(C6H6O3, 0.31), 123.1176(C9H14, 0.54), 121.1017(C9H12, 0.42), 119.0864(C9H10, 0.26), 111.0806(C7H10O, 0.99), 109.1017(C8H12, 1), 107.086(C8H10, 0.48), 97.1016(C7H12, 0.41), 95.0499(C6H6O3, 0.99), 83.0856(C6H10, 0.21) |
| 3.38 | 6467 | 485.3110 | $C_{26}H_{46}O_9$ [-$H_2O$+H]$^+$ | 485.3153(**C26H44O8**, 0.21), 467.2995(C26H42O7, 0.14), 428.2433(C22H35O8, 0.35), 410.2323(C22H33O7, 0.24), 383.2798(C22H38O5, 0.14), 323.258(**C20H34O3**, 0.33), 305.2484(C20H32O2, 0.57), 300.1599(C15H23O6, 0.13), 290.1752(C14H25O6, 0.12), 286.1448(C14H21O6, 0.2), 279.2324(C18H30O2, 0.13), 263.1659(C16H22O3, 0.12), 209.1904(C14H24O, 0.14), 167.0712(C9H10O3, 0.17), 149.0972(C10H12O, 0.22), 137.0961(C9H12O, 0.2), 135.0814(C9H10O, 0.24), 125.0974(C8H12O, 0.2), 121.1019(C9H12, 0.27), 111.0809(C7H10O, 0.13), 109.0653(C7H8O, 1), 107.0859(C8H10, 0.17), 97.1015(C7H12, 0.13), 95.0495(C5H5O, 0.44) |
| 3.29 | 6830 | 503.3227 | $C_{26}H_{46}O_9$ [+H]$^+$ | 485.3153(**C26H44O8**, 0.05), 467.3013(C26H42O7, 0.53), 365.2686(C22H36O4, 0.25), 363.2536(C22H34O4, 0.18), 323.2584(**C20H34O3**, 0.30), 305.2484(C20H32O2, 0.78), 277.2179(C18H28O2, 0.22), 235.1344(C14H18O3, 0.21), 209.1916(C14H24O, 0.2), 177.0554(C10H8O3, 0.24), 167.1444(C11H18O, 0.16), 161.0965(C11H12O, 0.24), 149.1334(C11H16, 0.25), 139.1112(C9H14O, 0.2), 135.0806(C9H10O, 0.16), 131.0859(C10H10, 0.15), 127.0398(C6H6O3, 0.19), 125.0968(C8H12O, 0.2), 123.1186(C9H14, 0.17), 121.1021(C9H12, 0.36), 109.0652(C7H8O, 1), 107.086(C8H10, 0.2), 95.0495(C6H6O, 0.51) |
| 3.24 | 7263 | 525.3031 | $C_{26}H_{46}O_9$ [+Na]$^+$ | 525.3019(**C26H46O9**, 1), 507.2928(C26H44O8, 0.07), 485.3114(C26H44O8, 0.02), 467.3006(C26H42O7, 0.2), 463.3039(C25H44O6, 0.08), 449.2903(C26H40O6, 0.02), 341.2686(**C20H36O4**, 0.02), 323.2581(C20H34O4, 0.13), 305.2483(C20H32O2, 0.37), 287.2374(C20H30O, 0.03), 209.1904(C14H24O, 0.08), 167.1432(C11H18O, 0.02), 149.1328(C11H16, 0.04), 139.1116(C9H14O, 0.01), 137.096(C9H12O, 0.01), 133.1016(C10H12, 0.01), 125.0967(C8H12O, 0.02), 123.1177(C9H14, 0.02), 121.1018(C9H12, 0.05), 119.086(C9H10, 0.02), 111.0804(C7H10O, 0.02), 109.0653(C7H8O, 0.18), 107.0859(C8H10, 0.01), 95.0494(C6H6O, 0.07) |
| 3.23 | 6827 | 503.3202 | $C_{26}H_{46}O_9$ [+H]$^+$ | 483.2936(**C26H42O8**, 0.16), 347.257(C22H34O3, 0.21), 321.2435(**C20H32O3**, 305.2474(C20H32O2, 0.33), 289.2529(C20H32O, 0.17), 137.0958(C9H12O, 0.16), 135.1183(C10H14, 0.21), 127.039(C6H6O3, 0.19), 123.1162(C9H14, 0.19), 121.1016(C9H12, 0.27), 0.23), 109.0657(C7H8O, 1), 95.0493(C6H6O3, 0.46) |

**Supplementary Table S10.** Continued.

| VIP | ID | *m/z* | Formula | Fragmentation spectrum *m/z*(formula, rel. intensity) |
|---|---|---|---|---|
| 3.14 | 6831 | 503.3231 | $C_{26}H_{46}O_{9}$ [+H]$^{+}$ | 467.3018(**C26H42O7**, 0.6), 449.2898(C26H40O6, 0.12), 365.2683(C22H36O4, 0.24), 363.2511(C22H34O4, 0.23), 341.2705(C20H36O4, 0.1), 323.2586(C20H34O3, 0.43), 305.2482(**C20H32O2**, 0.82), 287.2373(C20H30O, 0.15), 269.2278(C20H28, 0.06), 217.1963(C116H24, 0.03), 209.1904(C14H24O, 0.38), 195.1383(C12H18O2, 0.01), 193.1231(C12H16O2, 0.02), 191.1804(C14H22, 0.06), 177.1278(C12H16O, 0.01), 175.1119(C12H14O, 0.05), 167.1446(C11H18O, 0.08), 165.0917(C10H12O2, 0.07), 159.1174(C12H14, 0.04), 149.1327(C11H16, 0.16), 147.1176(C11H14, 0.03), 139.1123(C9H14O, 0.05), 137.0971(C9H12O, 0.06), 135.0811(C9H10O, 0.04), 125.0966(C8H12O, 0.07), 123.1176(C9H14, 0.1), 121.1019(C9H12, 0.2), 111.0808(C7H10O, 0.05), 109.0654(C7H8O, 1), 107.086(C8H10, 0.12), 97.1014(C7H12, 0.06), 95.0496(C6H6O, 0.21), 81.0711(C6H8, 0.04) |
| 3.12 | 6469 | 485.3117 | $C_{26}H_{46}O_{9}$ [-$H_2O$+H]$^{+}$ | 485.3167(**C26H44O8**, 0.01), 467.3006(C26H42O7, 0.29), 449.2904(C26H40O6, 0.09), 365.1699(C22H36O4, 0.13), 323.259(**C20H34O3**, 0.19), 305.2487(C20H32O2, 0.66), 295.2273(C18H30O3, 0.14), 287.2387(C20H30O, 0.14), 259.2059(C18H26O, 0.01), 217.1958(C16H24, 0.03), 209.1903(C14H24O, 0.09), 195.1382(C12H18O2, 0.12), 191.1808(C14H22, 0.05), 175.1491(C13H18, 0.01), 171.1025(C9H14O3, 0.01), 167.108(C10H14O2, 0.03), 165.0912(C10H12O2, 0.07), 163.113(C11H14O, 0.06), 161.1328(C12H16, 0.01), 159.118(C12H14, 7), 151.1121(C10H14O, 0.03), 149.1335(C11H16, 0.21), 147.1176(C11H14, 0.07), 137.0973(C9H12O, 0.07), 123.1172(C9H14O3, 0.06), 121.1016(C9H12, 0.22), 113.0969(C7H12O, 0.02), 111.081(C7H10O, 0.06), 109.0657(C7H8O, 0.66), 107.0859(C8H10, 0.06), 99.0808(C6H10O, 0.03), 97.1017(C7H12, 0.03), 95.0863(C7H10, 0.2), 83.0854(C6H10O, 0.02), 81.0704(C6H8, 0.07) |
| 3.12 | 7469 | 541.2978 | $C_{28}H_{44}O_{10}$ [+H]$^{+}$ | 541.2974(**C28H44O10**, 1), 539.284(C28H42O10, 0.17), 379.2472(**C22H34O5**, 0.1), 321.2428(C20H32O3, 0.07), 123.0797(C8H10O, 0.08), 109.0655(C7H8O, 0.08), 95.0492(C6H6O, 0.08) |
| 3.11 | 7092 | 519.3166 | $C_{28}H_{46}O_{10}$ [+H]$^{+}$ | 483.2964(**C26H42O8**, 0.31), 465.2857(C26H40O7, 0.39), 347.2558(C22H34O3, 0.84), 339.2545(C20H34O4, 0.22), 321.2428(**C20H32O3**, 0.38), 307.2638(C20H34O2, 0.37), 305.2487(C20H32O2, 0.31), 303.2336(C20H30O2, 0.3), 289.2534(C20H32O, 0.31), 209.1901(C14H24O, 0.17), 183.1025(C10H14O3, 0.17), 177.1639(C13H20, 0.14), 175.1124(C12H14O, 0.14), 165.0912(C10H12O2, 0.19), 149.1327(C11H16, 0.2), 145.0508(C6H8O4, 0.13), 139.1125(C9H14O, 0.14), 137.1336(C10H16, 0.18), 135.1184(C10H14, 0.22), 127.0759(C7H10O2, 0.16), 123.1174(C9H14, 0.35), 121.1016(C9H12, 0.25), 111.081(C7H10O2, 0.15), 109.0652(C7H8O, 0.84), 105.0704(C8H8, 0.21), 99.0454(C5H6O2, 0.14), 95.0497(C6H6O, 0.62) |
| 3.09 | 6097 | 467.3007 | $C_{26}H_{44}O_{8}$ [-$H_2O$+H]$^{+}$ | 467.2993(**C26H42O7**, 0.12), 449.287(C26H40O6, 0.03), 365.2697(C22H36O4, 0.05), 323.2591(C20H34O3, 0.08), 305.2481(**C20H32O2**, 0.49), 287.235(C20H30O, 0.09), 193.1222(C12H16O2, 0.03), 167.1068(C10H14O2, 0.02), 161.0958(C11H12O, 0.04), 151.1125(C10H14O, 0.02), 149.1329(C11H16, 0.1), 145.05(C6H8O4, 0.02), 137.0966(C9H12O, 0.03), 127.0395(C6H6O3, 0.12), 123.1176(C9H14, 0.08), 121.1017(C9H12O, 0.14), 109.0655(C7H8, 1), 107.0861(C8H10, 0.07), 103.0399(C4H6O3, 0.13), 97.0652(C6H8O, 0.15), 95.0494(C6H6O, 0.2), 85.0293(C4H4O2, 0.04), 81.0704(C6H8, 0.05), 69.7(C5H8, 0.05) |
| 3.08 | 7471 | 541.2993 | $C_{26}H_{46}O_{10}$ [+Na]$^{+}$ | 541.2989(C26H46O10, 1), 483.2954(C26H42O8, 0.05) |

**Supplementary Table S10.** Continued.

| VIP | ID | *m/z* | Formula | Fragmentation spectrum *m/z*(formula, rel. intensity) |
|---|---|---|---|---|
| 3.05 | 7091 | 519.3166 | $C_{30}H_{46}O_7$ [+H]$^+$ | 519.3276(C30H46O7, 0.17), 518.319(C30H45O7, 0.09), 517.3164(C30H44O7, 0.17), 515.2985(C30H42O7, 0.13), 503.2983(C29H42O7, 0.2), 447.3087(C27H42O5, 0.16), 399.275(C22H38O6, 0.1), 363.2518(C22H34O4, 0.1), 347.2564(C22H34O3, 0.18), 339.2537(C20H34O4, 0.18), 321.2439(C20H32O3, 0.45), 307.2644(C20H34O2, 0.22), 305.2483(C20H32O2, 0.1), 303.2332(C20H30O2, 0.16), 251.2007(C16H26O2, 0.09), 177.1285(C12H16O, 0.11), 167.1075(C10H14O2, 0.09), 161.1333(C12H16, 0.1), 159.1177(C12H14, 0.08), 149.0969(C10H12O, 0.16), 145.05(C6H8O4, 0.09), 139.1121(C9H14O, 0.13), 137.0977(C9H12O, 0.12), 135.0822(C9H10O, 0.14), 127.0768(C7H10O2, 0.12), 125.0966(C8H12O, 0.08), 123.1177(C9H14, 0.12), 121.1023(C9H12, 0.13), 111.0811(C7H10O2, 0.39), 109.0653(C7H8O, 0.36), 107.0861(C8H10, 0.12), 95.0493(C6H6O, 0.29) |
| 3.04 | 6466 | 485.3105 | $C_{26}H_{44}O_8$ [+H]$^+$ | 467.300330 (**C26H42O7**, 0.03), 305.2481(**C20H32O2**, 0.31), 149.1319(C11H16, 0.27), 137.1331(C10H16, 0.27), 123.1172(C9H14, 0.4), 121.1017(C9H12, 0.32), 111.0807(C7H10O, 0.3), 109.0654(C7H8O, 1), 107.0854(C8H10, 0.32), 97.1016(C7H12, 0.2542), 95.0499(C6H6O, 0.74) |
| 2.97 | 6776 | 501.3069 | $C_{26}H_{46}O_{10}$ [-$H_2O$+H]$^+$ | 501.3069(C26H44O9, 0), 483.2950 (**C26H42O8**, 0.07), 321.2426(**C20H32O3**, 0.18), 307.2648(C20H34O2, 0.4), 305.2478(C20H32O2, 0.24), 293.2126(C18H28O3, 0.77), 278.2214(C18H29O2, 0.14), 277.2176(C18H28O2, 0.62), 171.1181(C13H14, 0.13), 165.0916(C10H12O2, 0.15), 161.1333(C12H16, 0.15), 159.1176(C12H14, 0.15), 151.1133(C10H14O, 0.27), 149.1332(C11H16, 0.28), 147.117(C11H14, 0.19), 145.1017(C11H12, 0.15), 137.0965(C9H12O, 0.24), 135.1179(C10H114, 0.32), 133.1021(C10H12, 0.14), 123.1172(C9H14, 0.24), 121.102(C9H12, 0.33), 119.0859(C9H10, 0.14), 111.0815(C7H10O, 1), 107.0862(C8H10, 0.37), 105.0709(C8H8, 0.17), 97.1015(C7H12, 0.16), 94.0741(C7H9, 0.64), 81.0707(C6H8, 0.13) |
| 2.94 | 6156 | 469.3159 | $C_{26}H_{44}O_7$ [+H]$^+$ | 450.297590(C**26H42O6**, 0.05), 289.2527(**C20H32O**, 0.22), 123.1174(C9H14, 0.23), 121.1016(C9H12, 0.24), 109.0657(C7H8O, 0.28), 93.07(C7H8O, 1) |
| 2.94 | 7472 | 541.2999 | $C_{26}H_{46}O_{10}$ [+Na]$^+$ | 541.2984(C26H46O10, 1), 539.2816(C26H44O10, 0.2), 537.2634(C26H42O10, 0.08), 505.2775 (**C26H42O3**, 0.03), 321.2425(**C20H32O3**, 0.09), 123.1177(C9H14, 0.08), 109.065(C7H8O, 0.13), 95.0496(C6H6O, 0.1) |
| 2.91 | 7266 | 525.3039 | $C_{28}H_{44}O_9$ [+H]$^+$ | 525.3032(**C28H44O9**, 1), 523.2892(C28H42O9, 0.01), 507.2931(C28H42O8, 0.08), 467.2997(C26H42O7, 0.13), 463.3044(C27H42O6, 0.06), 365.2697(C22H36O4, 0.02), 363.2527(**C22H34O4**, 0.30), 323.2577(C20H34O3, 0.06), 305.249(C20H32O2, 0.17), 209.1904(C14H24O, 0.02), 149.1333(C11H16, 0.03), 121.1015(C9H12, 0.01), 109.0652(C7H8O, 0.11), 107.0859(C8H10, 0.02), 95.0494(C6H6O, 0.03) |

#### Supplementary Figures

A)

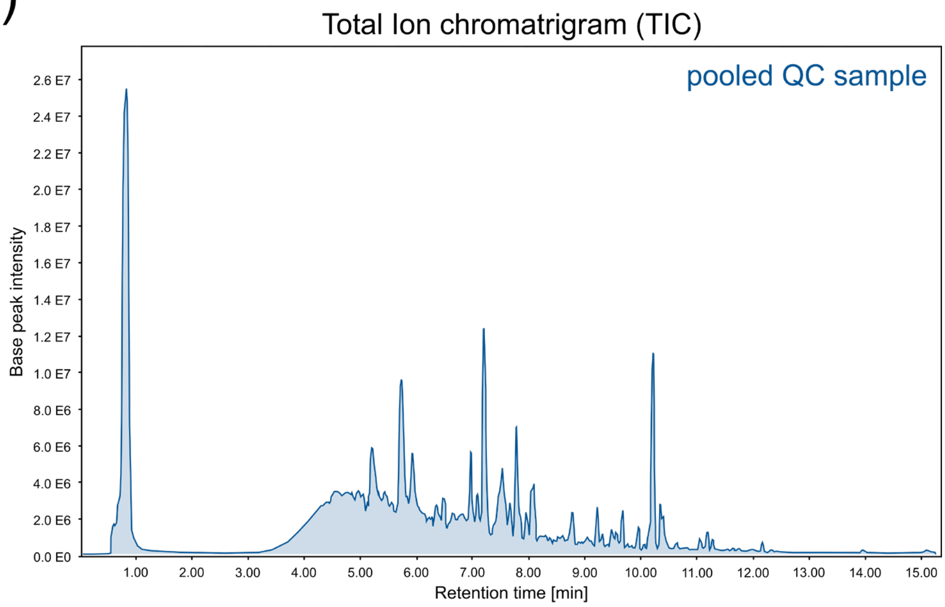

B)

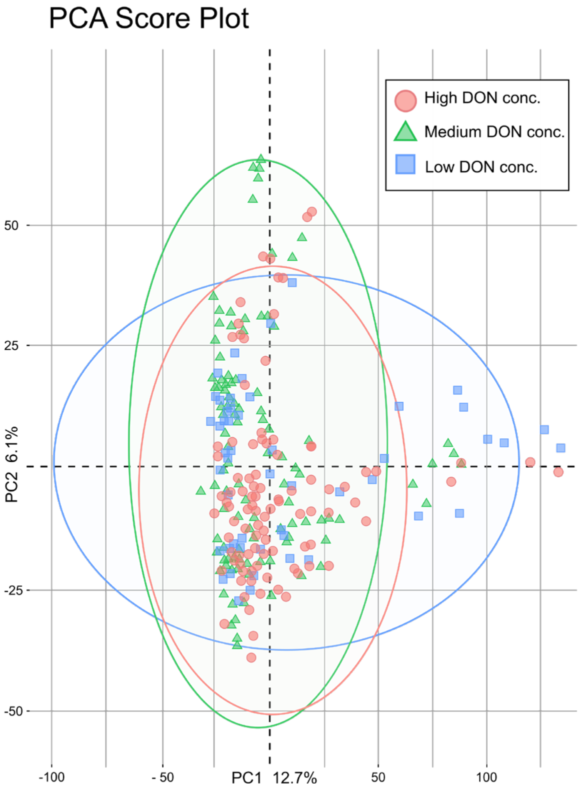

**Supplementary Figure S1.** Total ion chromatogram of the pooled QC sample in ESI(+)-UPLC-TOF-MS (A) and PCA of the processed data matrix (B). The TIC shows the base peaks, the PCA is colored by the samples' DON concentration (high: > 1,240 mg/kg, medium: > 620 mg/kg low: < 10 mg/kg) and confidence ellipses (95%) are plotted in the respective color.

### A) Linear correlation of features most significantly positively related to DON-toxin load

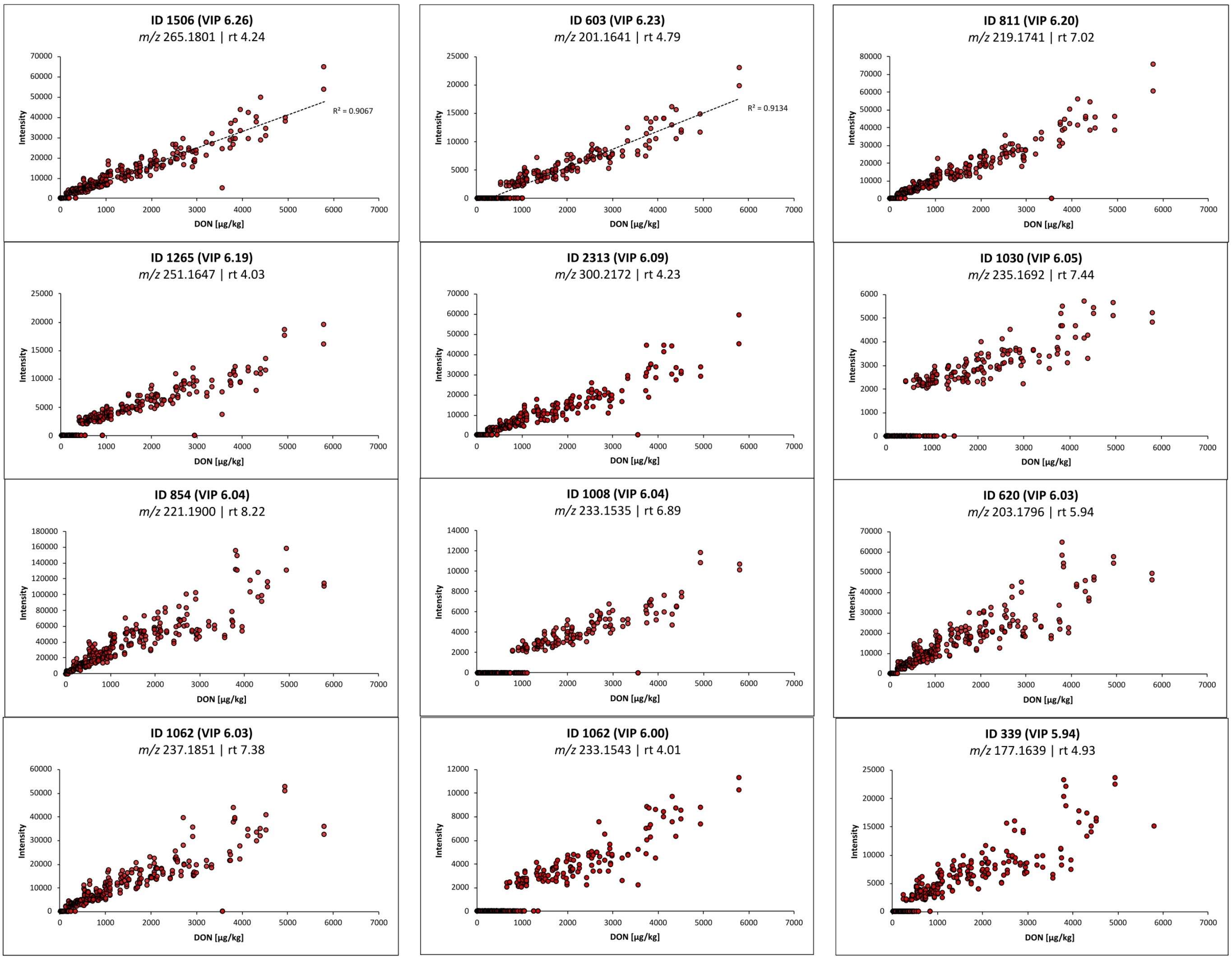

### B) Correlation of features most significantly negatively related to DON-toxin load

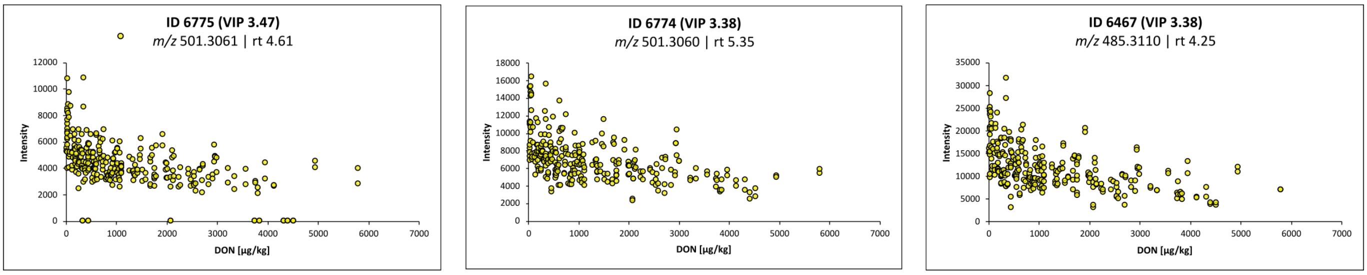

**Supplementary Figure S2.** The intensities of most significant positively (A) and negatively (B) correlating features against the measured DON-concentration in the respective samples and replicates.

#### A) Regression model training and testing

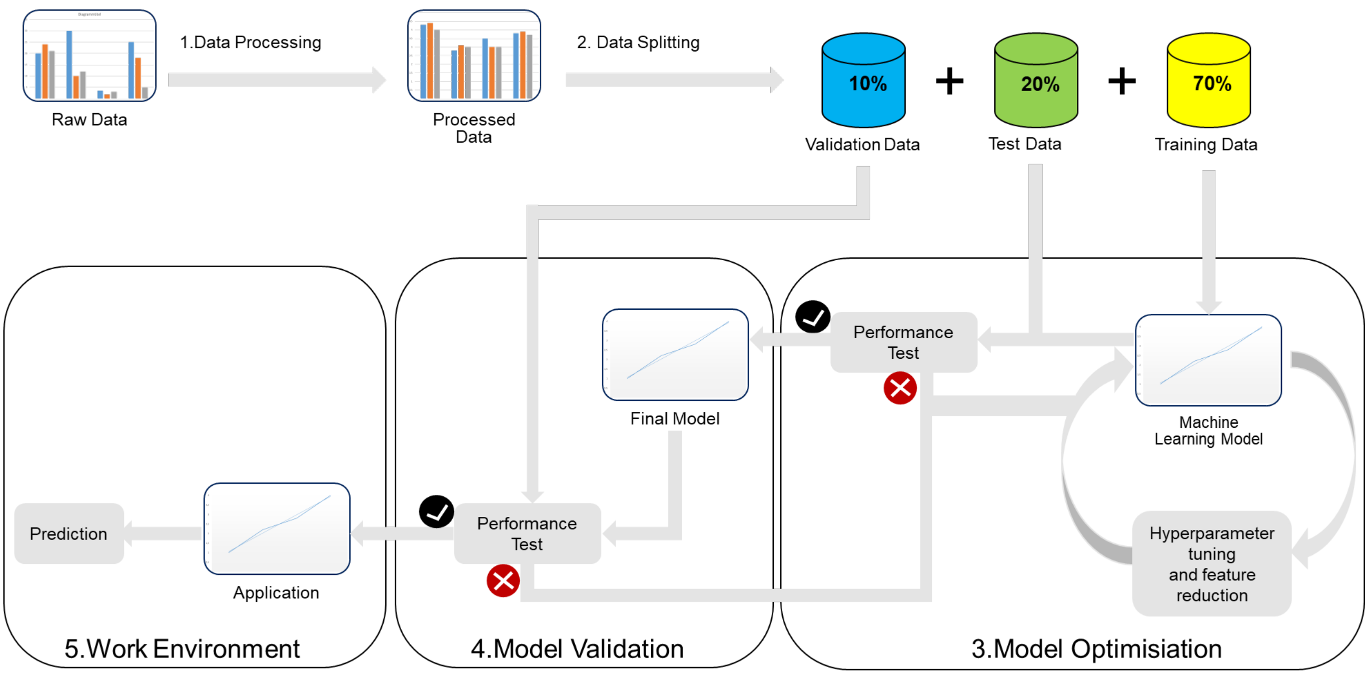

#### B) Linear, LASSO, and PLS regression model coefficients of determination

| Model | Model details | | |
|---|---|---|---|
| | Data structure | Test $R^2$ | Validation $R^2$ |
| Linear regression | z-score normalized | 0.98 | 0.97 |
| | min-max normalized | 0.96 | 0.98 |
| LASSO regression | z-score normalized | 0.98 | 0.97 |
| | min-max normalized | 0.98 | 0.98 |
| PLS regression | z-score normalized | 0.96 | 0.95 |
| | min-max normalized | 0.96 | 0.98 |

**Supplementary Figure S3.** Training and testing concept (A) and the coefficients of determination values (B) of linear, LASSO, and PLS model regressions predicting the DON toxin load based on metabolomic data.
